# Leading and lagging strand abasic sites differentially affect vertebrate replisome progression but involve analogous bypass mechanisms

**DOI:** 10.1101/2025.01.09.632187

**Authors:** Matthew T. Cranford, Steven N Dahmen, David Cortez, James M. Dewar

**Affiliations:** Department of Biochemistry, Vanderbilt University School of Medicine, Nashville, TN, USA

## Abstract

Abasic sites are one of the most frequent forms of DNA damage that interfere with DNA replication. However, abasic sites exhibit complex effects because they can be processed into other types of DNA damage. Thus, it remains poorly understood how abasic sites affect replisome progression, which replication-coupled repair pathways they elicit, and whether this is affected by the template strand that is damaged. Using *Xenopus* egg extracts, we developed an approach to analyze replication of DNA containing a site-specific, stable abasic site on the leading or lagging strand template. We show that abasic sites robustly stall synthesis of nascent DNA strands but exert different effects when encountered on the leading or lagging strand template. At a leading strand AP site, replisomes stall ∼100 bp from the lesion until it is bypassed or a converging fork triggers termination. At a lagging strand abasic site, replisome progression is unaffected and lagging strands are reprimed downstream, generating a post-replicative gap, which is then bypassed. Despite different effects on replisome progression, both leading and lagging strand abasic sites rely on translesion DNA synthesis for bypass. Our results detail similarities and differences between how leading and lagging strand AP sites affect vertebrate DNA replication.

## Introduction

Approximately 15,000 abasic sites (apurinic/apyrimidinic sites; abbreviated ‘AP sites’) form per day in each human cell through spontaneous base loss or excision of damaged bases by DNA glycosylases (1–5). Recent measurements suggest a steady state level of one AP site per 1.25-2.5×10^6^ nucleotides (6). This frequency is increased dramatically by many environmental genotoxins including those derived from foods and industrial processes, as well as ionizing and UV radiation (7). The base excision repair pathway rapidly repairs AP sites, but their high frequency and preferential formation within single-stranded DNA (8,9), means that they are inevitably encountered during DNA replication (10). AP sites are genotoxic (7) and cytotoxic (11). Furthermore, AP sites can be converted into single-stranded DNA breaks (SSBs) (12,13), DNA-protein cross-links (DPCs) (14,15), and DNA interstrand cross-links (ICLs) (16). Because AP sites are one of the most frequent forms of DNA damage experienced by cells (1–5), it is crucial to understand how they affect DNA replication.

At eukaryotic replication forks, the replicative helicase translocates on the leading strand template and forms a macromolecular ‘replisome’ complex with DNA polymerases and other replication proteins, to perform DNA synthesis (17–20). AP sites stall some, but not all, helicases (21–27), and readily stall DNA polymerases (2,4,5) but it is less clear how AP sites affect helicase and DNA polymerase activities of the eukaryotic replisome. AP sites on the lagging strand template are not expected to block replisome progression because the lesion is on the non-translocating strand and lagging strand synthesis is naturally discontinuous (28,29), which allows for re-initiation of lagging strand synthesis downstream of DNA damage (30,31). AP sites on the leading template strand should not block the replicative helicase because they are smaller than the unmodified bases that are normally accommodated within the central channel (18). However, full replicative helicase activity requires coupled DNA synthesis by the leading strand polymerase (32–34), so replisome progression may be impaired by a leading strand AP site. Accordingly, reconstituted yeast replisomes progress more slowly after encountering a leading strand AP site, as for other base modifications (35). However, replisome stalling was not detected, and it is unclear how close to the AP site replisomes slow (35). An analysis of AP site replication in *Xenopus* egg extracts detected stalling of nascent strands but did examine replisome stalling (36) and thus did not address this point. Overall, it is unknown whether the eukaryotic replisome stalls at an AP site and, if so, how close to the lesion this occurs.

After a replicative DNA polymerase stalls at an AP site, a DNA damage tolerance mechanism may be needed to complete DNA synthesis. Mutagenesis studies in yeast suggest that completion of synthesis primarily involves error-prone translesion DNA synthesis (TLS) polymerases (37–40), which are sufficient to bypass AP sites in biochemical reconstitution experiments (41–44). TLS-independent bypass of DNA lesions is thought to occur by fork reversal or template switching (30,31), and at least one of these pathways was suggested to support AP site bypass in *Xenopus* egg extracts (36). Accordingly, the homologous recombination protein RAD51 and its regulators, which participate in both template switching and fork reversal (30,31), can bind single-stranded DNA containing AP sites (45,46). Replicative DNA polymerases can also bypass AP sites at reduced efficiency (37–40,44,47,48), which represents an additional TLS-independent bypass mechanism. Bypass of DNA lesions can occur through both post-replicative and on-the-fly mechanisms (49–55), but the frequency of these outcomes in the context of AP sites remains unclear. Thus, multiple mechanisms are implicated in synthesis past AP sites, but it is unclear to what extent they are deployed during AP site replication in vertebrates and to what extent they occur close to the fork or at post-replicative gaps.

To understand how AP sites on leading or lagging template strands affect DNA replication, we utilized *Xenopus* egg extracts (56) that support replication coupled DNA repair (36,57,58) using a complex set of vertebrate nuclear proteins (59,60). This approach allowed us to replicate custom DNA templates that were engineered to contain a site-specific AP site on either the leading or lagging strand template. Our study provides a high temporal- and spatial-resolution view of AP site replication, which shows that AP sites robustly stall DNA synthesis but are ultimately bypassed. Surprisingly, a leading strand template AP site rapidly stalls the replisome with lagging strands located ∼100 nucleotides beyond the AP site until restart of DNA synthesis or termination by a converging fork. This contrasts to the continued unwinding and lagging strand synthesis predicted by other studies (35). Replisomes are not impeded by lagging strand template AP sites, and nascent lagging strands reprime downstream of the damage, as expected (61). Despite the stark differences between replisome behavior, the mechanism of bypass in both cases requires TLS. Furthermore, blocking leading strand bypass revealed that there is minimal uncoupling of leading and lagging strand synthesis, while blocking lagging strand bypass did not affect post-replicative gap formation. Overall, our data reveal that leading and lagging template strand abasic sites exert remarkably different effects on replisome progression despite using analogous bypass mechanisms.

## Materials and Methods

### Construction and validation of AP site plasmids

A plasmid containing a 50x lacO array (pJD161) (62) was linearized with PsiI (New England Biolabs). DNA oligonucleotides oMC30 and oMC31 (Supplemental Table 1) were annealed to form a blunt 73 base pair (bp) duplex (oMC30/31, Fig. S1A) which was ligated into the linearized plasmid. Plasmids were sequence-verified to isolate clones containing the oMC30/31 insert in either the forward (pMC9) or reverse (pMC10) orientation. This novel insert contains tandem Nt.BbvCI nicking sites spaced 63 bp apart and serves as a receptor sequence for modified oligonucleotides (Supplemental Table 1) used to generate AP site plasmids. Thus, pMC9 contains the Nt.BbvCI nicking sites in the top strand to allow for modification of the lagging strand template, and pMC10 contains the Nt.BbvCI nicking sites in the bottom strand to allow for modification of the leading strand template (Fig. S1A). To generate a plasmid without the 50x lacO array, pMC10 was digested with BsrGI and BsiWI (New England Biolabs). The target fragment without the 50x lacO array was gel purified, re-ligated and sequence-verified to validate generation of a plasmid containing the receptor sequence insert but without the lacO array (pMC14).

To construct modified plasmids, pMC9, pMC10, or pMC14 was nicked with Nt.BbvCI (New England Biolabs). Nicked products were then purified using a PCR purification kit (Qiagen). Modified oligonucleotides (Supplemental Table 1) were annealed into the nicked plasmid by mixing >100-fold molar excess of indicated oligonucleotides with nicked plasmids in 10 mM Tris-HCl. Annealed products were ligated with T4 DNA ligase (New England Biolabs) then treated with T5 exonuclease (New England Biolabs) to digest excess unligated plasmids and oligonucleotides, similar to previously described (63). Ligated products were then purified using a PCR purification kit followed by 3 rounds of buffer exchange with 500 μl of 10 mM Tris-HCl (pH 8) through a 100 kDa centrifugal filter (Amicon). Modified plasmids were diluted with 10 mM Tris-HCl (pH 8) and stored at a final concentration of 150 ng/μl.

The position of the dUracil modification is incorporated within a MscI restriction site (Fig. S1A). Therefore, construction of modified plasmids was validated by screening for resistance to digestion by MscI. To validate modified plasmids, 300 ng of plasmids were digested with 5 units of XmnI with or without 5 units of MscI in 1x rCutsmart buffer (New England Biolabs) in a reaction volume of 10 μl at 37°C. Digest products were brought to 1x DNA loading buffer (3.3 mM Tris-HCl pH 8, 0.017% SDS, 11 mM EDTA, 0.015% Bromophenol Blue, 2.5% Ficol) and resolved through a 0.8% TBE-agarose gel (0.3 μg/mL ethidium bromide). All modified plasmids were successfully linearized by XmnI, as expected. However, only the plasmids modified with the undamaged oligonucleotide sequence were sensitive to digestion by MscI, whereas all dUracil-modified plasmids were fully resistant to MscI. An example validation is shown for different modifications of pMC14 (Fig. S1C).

To generate AP site plasmids, 225 ng of modified plasmids were treated with 3 units of uracil-DNA glycosylase (UDG, New England Biolabs) in 1x UDG buffer in a final volume of 3 μl at 37°C for 30 minutes. This approach was used to generate AP site plasmids to validate AP site generation, for stability and retention of AP sites in extracts (see below), and in replication assays (see below). Generation of AP sites was screened for by digesting 1 μl of UDG-treated plasmids with AP Endonuclease (APE1) in 1x NEB 4 buffer (New England Biolabs) in a reaction volume of 10 μl at 37°C for 30 minutes. Products were brought to 1x DNA loading buffer and resolved through a 0.8% TBE-agarose gel (0.3 μg/mL ethidium bromide). All modified plasmids demonstrated partial sensitivity to APE1 in the absence of UDG, but only dUracil-modified plasmids were fully sensitive to APE1 after treatment with UDG. An example validation of AP site generation is shown (Fig. S1D).

### Expression and purification of LacR

Biotinylated LacR was expressed in *E. coli* and purified as described previously (64,65).

### Preparation of *Xenopus* egg extracts

High-speed supernatant (HSS) and nucleoplasmic extract (NPE) *Xenopus* egg extracts were prepared from *Xenopus* laevis (Nasco) as previously described (66). Animal protocols were approved by Vanderbilt Division of Animal Care (DAC) and Institutional Animal Care and Use committee (IACUC). HSS was activated by supplementing with nocodazole (3 ng/μl) and ATP regenerating system (ARS; 20 mM phosphocreatine, 2 mM ATP, and 5 ng/μl creatine phosphokinase) and centrifuged at 21,130 RCF for 5 minutes. The activated HSS was harvested and used for licensing plasmids. NPE was supplemented with ARS, DTT (2 mM), and diluted to a final concentration of 50% (v/v%) in 1x egg lysis buffer (ELB; 250 mM sucrose, 2.5 mM MgCl2, 50 mM KCl, 10 mM HEPES pH 7.7).

### Stability and retention of AP sites in extracts

Modified plasmids were treated with UDG to generate AP sites as described above. Plasmids were mixed with LacR (at a final concentration of 56.3 ng/μl of UDG-treated plasmids and 8.16 μM of LacR) and incubated at room temperature for 60-90 minutes to form the LacR array. To test stability and retention of AP sites in different extracts, 1 volume of LacR-bound plasmids was mixed with 4 volumes of 1x ELB (negative control), activated HSS, or non-licensing extracts (i.e. 50% NPE (v/v%) mixed with activated HSS in 2:1 ratio) and incubated at room temperature for 30 minutes. Samples were stopped with 10 volumes of Extraction Stop buffer (50 mM Tris-HCl pH 7.5, 25 mM EDTA, 0.5% SDS (w/v%)) and processed for analysis (see below).

### Replication assay in extracts

Modified plasmids were treated with UDG and mixed with LacR as outlined above. To license plasmids, UDG-treated and LacR-bound plasmids were mixed with activated HSS in a 1:4 ratio then incubated at room temperature for 30 minutes. 50% NPE (v/v%) was prepared as above and supplemented with [α-32P]dATP to label nascent DNA replication products upon initiate of replication. To initiate replication, 2 volumes of 50% NPE (v/v%) were mixed with 1 volume of licensing mix. Samples were taken at indicated time points by mixing with 10 volumes of Extraction Stop buffer and processed for analysis (see below). Where indicated, NPE alone was supplemented with etoposide (Etop; Sigma) to a final concentration of 100 μM in the reaction, and ubiquitin vinyl sulfone (UbVS; R&D Systems) was added to both the licensing mix and 50% NPE (v/v%) at a final concentration of 20 μM as previously described (64).

### Sample Processing and Analysis

In all experiments, samples were stopped with 10 volumes of Extraction Stop buffer, then treated with RNAse (0.3 mg/mL final) at 37°C for 30 minutes followed by Proteinase K (0.7 mg/mL final) at 37°C for 1 hour or at room temperature overnight, unless indicated otherwise (Fig. S2I).

To test for stability of AP site plasmids in extracts (Fig. S1F), processed samples were diluted in 1x DNA loading buffer and resolved by 0.8% agarose gel (w/v%) in 1xTBE and 0.3 μg/ml ethidium bromide. Gels were washed in 1x TBE for 30 minutes then stained in 1x SYBR Gold (Invitrogen) in 1x TBE for 30 minutes. To further test for retention of AP sites in extracts, samples were purified using Monarch PCR & DNA Cleanup Kit (New England Biolabs) and eluted in 10 mM Tris-HCl (pH 8). Purified samples were then digested with 0.2 units/μl XmnI and 0.05 units/μl MscI in 1x rCutsmart buffer for 30 minutes, diluted to 1x DNA Loading buffer and resolved by a 0.8% agarose gel (w/v%) in 1xTBE and 0.3 μg/ml ethidium bromide. Gels were washed in 1x TBE for 30 minutes then stained in 1x SYBR Gold in 1x TBE for 30 minutes. Retention of the AP site was measured by resistance of the 3 kb fragment to digestion by MscI (Fig. S1G-H).

For replication assays, processed samples were diluted to 1x DNA loading buffer, resolved by 1% agarose gel (w/v%), and the gel was vacuum dried in a gel dryer and visualized by autoradiography. To detect DPCs, processed samples (± Proteinase K) were diluted 10-fold in Replication Stop buffer (80 mM Tris-HCl (pH 8), 8 mM EDTA, 5% SDS, 10% Ficoll, 0.13% phosphoric acid, 0.2% bromophenol blue) and resolved by native agarose gel, vacuum dried in a gel dryer and visualized by autoradiography. For 2D gel analyses, processed samples were diluted to 1x DNA loading buffer and resolved in the first dimension on a 0.4% agarose gel (w/v%). Gel slices containing products of desired size were stained in 0.3 μg/mL ethidium bromide (in 1x TBE), rotated 90°, and cast in a 1.2% agarose gel (w/v%) supplemented with 0.3 μg/mL ethidium bromide. Products were resolved in the second dimension at 4°C then vacuum dried in a gel dryer and visualized by autoradiography. For denaturing gel analyses, replication products were purified using a Monarch PCR and DNA Cleanup kit and eluted in 10 mM Tris-HCl (pH 8). Purified replication products were digested with 0.2 units/μl of the indicated restriction enzymes in 1x rCutsmart buffer at 37°C for 30 minutes. Digests were stopped by addition of EDTA (30 mM final) then mixed into 1x alkaline DNA sample buffer (50 mM NaOH, 1 mM EDTA, 3% Ficol (w/v%), 0.026% bromocresol green (w/v%), 0.043% xylene cyanol (w/v%)). Samples were resolved through a 1.5% agarose gel (w/v%) in 1x alkaline agarose buffer (50 mM NaOH, 1 mM EDTA). The gel was neutralized in 7% trichloroacetic acid solution, washed in 1x TE, then vacuum dried in a gel dryer and visualized by autoradiography. For sequencing gels, samples were digested as above then mixed with an equal volume of Gel Loading Buffer II (Invitrogen) and resolved through a 6% urea-polyacrylamide sequencing gel. The gel was dried under vacuum and visualized by autoradiography.

Quantification of stalled and full-length replication products was performed using ImageLab v6.1 (BioRad). Briefly, Volume Tools was used to acquire measurements for total lane signal and signal of indicated product bands. Background signal from an adjacent region of the lane was subtracted from respective measurements. In cases where background signal of a measurement was inconsistent across a time course (due to synthesis of additional replication intermediates of similar size), average background signals (taken from above and below respective measurements) were subtracted (e.g. Stalled Lagging in Fig. 4B-C, Fig. S11E-G, Fig. S12 E-G). Quantification of indicated product bands is expressed as a percentage of total lane signal unless otherwise indicated. Lane profiles were generated using ImageQuantTL v8.2 (Cytiva). 1D gel analysis was used to generate lanes. Lane profiles (i.e. signal as a function of Rf) were exported to Excel. Lane signals were normalized to the signal of the opposing fork (e.g. see Fig. S5A,E). The x-axis was plotted in reverse order to represent Relative Product Size from small to large. All plots were composed using Prism v10.4.0 (GraphPad).

### Immunodepletions

Antibodies against Rev1 (Rev1-N and Rev1-C) were described previously (67). Antibodies against Polκ were re-generated using a peptide antigen as previously described (68). All immunodepletions were performed as previously described (69). Briefly, Protein A-coupled magnetic Dynabeads (Invitrogen) were washed and equilibrated in 1x PBS supplemented with 0.02% Tween. For every 1 μl of equilibrated beads, 0.75 μl of antibody serum (for Rev1-N and Rev1-C) or 0.5 μl of affinity purified antibody (1 mg/mL, for Polκ) was added and incubated at 4°C with end-over-end rotation overnight. Antibody-bound beads were washed twice with 1x PBS supplemented with 0.02% Tween, twice with 1x ELB supplemented with 500 mM NaCl and 0.02% Tween, once with 1x ELB supplemented with 0.02% Tween, then resuspended in 1x ELB. For each round of immunodepletion, 8.6 μl of antibody-bound beads were mixed with 20 μl of activated HSS, and 7.8 μl of antibody-bound beads were mixed with 18 μl of 50% NPE (v/v%) then incubated with end-over-end rotation at room temperature for 20 minutes. To deplete Rev1, HSS was depleted with 1 round of anti-Rev1-N beads followed by 1 round of anti-Rev1-C beads, and NPE was depleted by 2 rounds of anti-Rev1-N beads followed by 1 round of anti-Rev1-C beads. To deplete Polκ, HSS was depleted with 1 round of anti-Polκ beads and NPE was depleted by 3 rounds of anti-Polκ beads. Depleted extracts were harvested and used for replication assays as described above. Mock and depleted NPE were analyzed by western blotting to verify depletion of target proteins.

## Results

### Generation of a site-specific AP site plasmid

We wanted to investigate the effect of a site-specific AP site on DNA replication in *Xenopus* egg extracts independent of other AP-derived DNA damage. DNA SSBs are protected from repair in *Xenopus* egg extracts when flanked by DNA binding proteins (70). Therefore, we tested whether flanking an AP site with DNA-bound *lac* repressor (LacR) would protect it from modification (Fig. S1A). To this end, an oligonucleotide that contained deoxyuracil (dUracil) flanked by *lac* operator (*lacO*) sequences was annealed and ligated into a gapped recipient plasmid (Fig. S1B-C, as in (65)). The dUracil-containing plasmid (pdUracil) was then treated with Uracil DNA glycosylase (UDG) to convert the dUracil to an AP site (Fig. S1D). AP site insertion was highly efficient as evidenced by sensitivity of the AP site plasmid to AP endonuclease (APE1; Fig. S1D, lane 6). Upon incubation with *Xenopus* egg extracts, the AP site plasmid rapidly lost sensitivity to APE1 (Fig. S1E, lanes 1-4) indicating that the AP site was repaired, as expected (36). In contrast, LacR-bound plasmids were predominantly nicked (Fig. S1E, lanes 5-8), which indicated that repair was inhibited but the AP site was susceptible to nucleases that target AP sites, most likely APE1 (13). To confer nuclease resistance to the AP site plasmid, we introduced phosphorothioate linkages adjacent to the AP site. The phosphorothioate modification blocks APE1 activity (71). We then assessed repair of the AP site by restriction digestion with MscI, which is inhibited by the presence of an AP site. Using this approach, we found that the combination of LacR binding and phosphorothioate linkages stabilized essentially all AP sites (Fig. S1F-H). Thus, a phosphorothioate-containing and LacR-bound AP site was stable in *Xenopus* egg extracts and is hereafter referred to as a stable AP site.

### AP sites robustly stall DNA synthesis

To characterize how stable AP sites affect DNA replication, plasmid DNA templates that contained a stable AP site or the undamaged control sequence were replicated in *Xenopus* egg extracts (Fig. 1A). Replication of control DNA initially gave rise to θ structures (θs) due to the presence of replication forks on the DNA (Fig. 1A and Fig. 1B, lanes 1-2), as expected (64,65). By 30 minutes, almost all products were converted to supercoiled and nicked circular monomers (scCMs and nCMs, Fig. 1B, lanes 3-6, Fig. 1C, Fig. S2A) that are the final products of replication (56). Replication of a stable AP site plasmid similarly led to formation and resolution of θs with only a slight delay compared to the undamaged plasmid (Fig. 1B, lanes 2,8, Fig. S2B-C). The lack of persistent θs indicates that the AP sites did not form ICLs, which would have caused prolonged stalling of converging forks (72,73). We observed a slight enrichment of catenanes (Fig. 1B lanes 1-2,7-8, Fig. S2D-E), suggesting the AP site may slightly inhibit decatenation during termination of DNA replication (64). We also observed minimal formation of σ structures (Fig. S2F-H), which arise from replication of a SSB (70), and did not observe appreciable DPC formation (Fig. S2G). These data show that the stable AP site underwent DNA replication without evidence of substantial conversion to ICLs, SSBs, or DPCs.

**Figure 1:**
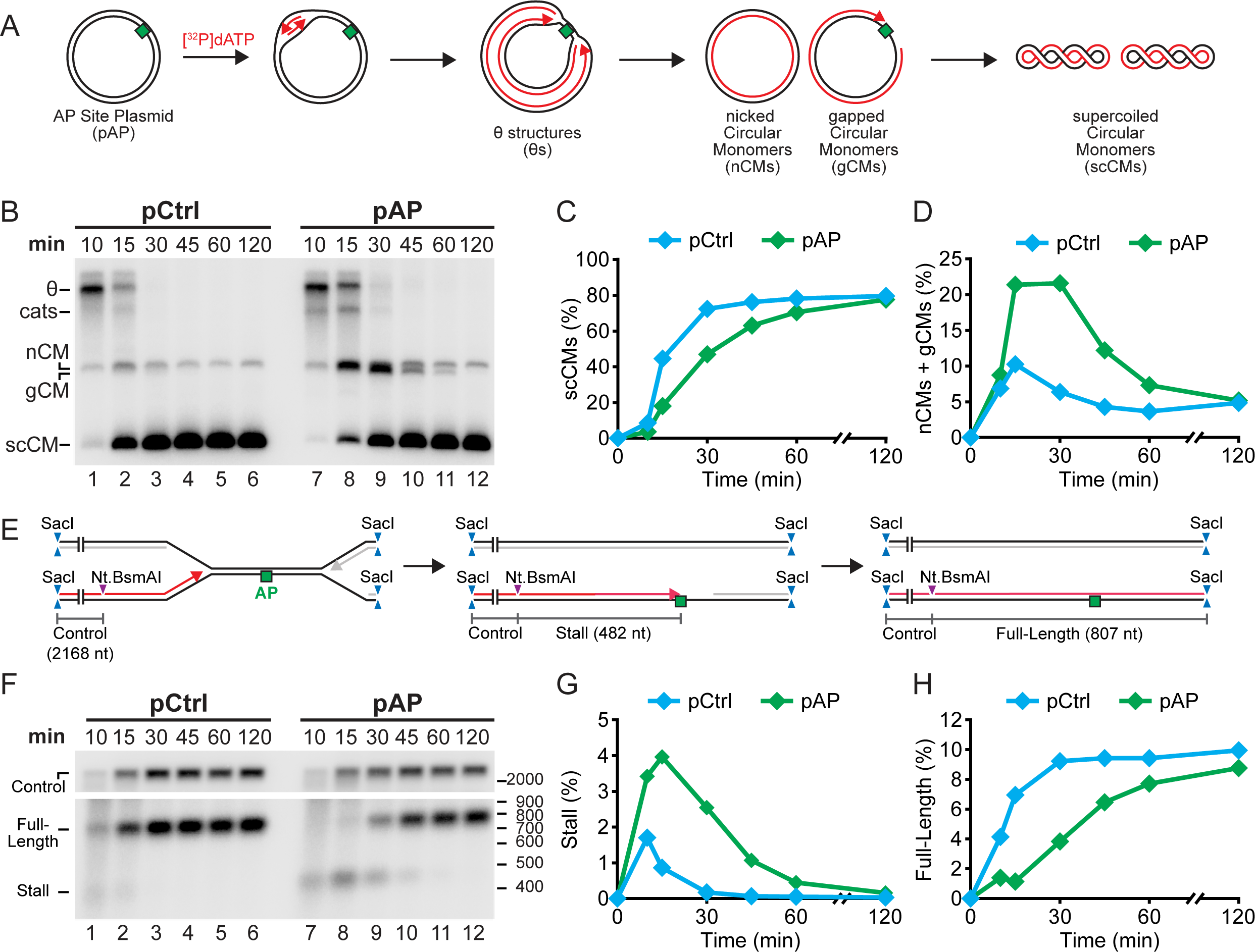
A stable AP site causes site-specific replication stalling in *Xenopus* egg extracts. A) Plasmid DNA harboring a stable AP site (pAP) was replicated in *Xenopus* egg extracts alongside control (pCtrl) DNA. [α-32P]-dATP was added to label newly-synthesized DNA strands. B) Products from (A) were resolved on a native agarose gel and visualized by autoradiography. C) Quantification of supercoiled circular monomers (scCMs) from (B). An independent experimental replicate is shown in Fig. S2A. D) Quantification of nicked and gapped circular monomers (nCM and gCM, respectively) from (B). An independent experimental replicate is shown in Fig. S2J. E) Purified replication intermediates from (A) were restriction-digested with SacI and Nt.BsmAI to distinguish stalled and nascent strands from full-length nascent strands. A fragment arising from nascent strands distal to the AP site serves as an internal loading control. F) Nascent DNA strands from (E) were separated on an alkaline denaturing agarose gel then visualized by autoradiography. Additional nascent strand products generated by this analysis are shown in Fig. S3C-F. G) Quantification of stall intermediates from (F). An independent experimental replicate is shown in Fig. S3A. H) Quantification of full-length products from (F). An independent experimental replicate is shown in Fig. S3G.

In contrast to the control, the AP site delayed formation of scCMs by approximately 15 minutes (Fig. 1B-C, Fig. S2A). During this time, nicked and gapped circular monomers accumulated (Fig. 1B, lanes 8-11, Fig. 1D, Fig. S2J). These nicked and gapped products arise from incomplete synthesis of the daughter DNA strands (Fig. 1A). Thus, the AP site interfered with completion of DNA synthesis. Importantly, replication of the AP site plasmid ultimately generated supercoiled products that were indistinguishable from the control (Fig. 1B lanes 6,12, Fig. 1C), indicating that the AP site was not a permanent block to DNA synthesis. The timing of scCM formation was consistent with efficient stalling of half of all nascent DNA strands by the AP site (Fig. S2K) (58). Altogether, these data establish that replication of the stable AP site delays complete synthesis of daughter DNA strands on one template DNA strand, suggesting strand-specific stalling by the AP site.

To directly monitor synthesis of nascent strands in the vicinity of the AP site, purified replication intermediates were restriction-digested (Fig. 1E) then separated using a denaturing agarose gel (Fig. 1F). This approach allowed us to separate daughter strands synthesized across the AP site from those synthesized on the undamaged strand (Fig. 1E). Replication of the AP site resulted in a prominent stall intermediate (Fig. 1E, Fig. 1F lanes 7-9, Fig. 1G, Fig. S3A) which declined over approximately 30 minutes (Fig. 1F, lanes 7-10, Fig. 1G, Fig. S3A). Replication of control DNA led to a less prominent stall intermediate that declined over approximately 5 minutes (Fig. 1F lanes 1-2, Fig. 1G, Fig. S3A). These less-prominent stall products were likely due to transient stalling at the LacR-bound *lacO* sites that flank the position of the AP site modification (Fig. S1A, hereafter referred to as ‘transient stalling’). The substantially increased abundance and persistence of the AP site stall product compared to control DNA (Fig. 1G, Fig. S3A) demonstrated that most stalling was attributable to the AP site. Importantly, nascent strands adjacent to the AP site region formed at a similar rate during replication of both AP site and control plasmids (Fig. 1F lanes 2 and 8, Fig. S3B), and were synthesized on the undamaged strand at a similar rate in both conditions (Fig. S3C-F). Thus, impaired synthesis was specific to the damaged strand within the vicinity of the AP site. Loss of the stall intermediate corresponded to the appearance of full-length strands (Fig. 1E, Fig. 1F lanes 9-12, Fig. 1H, Fig. S3G), which were delayed by approximately 30 minutes compared to control DNA (Fig. 1F lanes 2-3,9-10, Fig. 1H, Fig. S3G). Therefore, the AP site delayed, but did not block, replication of the damaged DNA strand. Overall, these data show that nascent DNA strands initially stall at the AP site but ultimately restart and complete DNA synthesis.

AP sites are expected to stall DNA polymerases, but not the replicative helicase, resulting in nascent DNA strands stalled one nucleotide before the damaged base (36). To test whether this was the case in our system, purified replication intermediates were restriction-digested (Fig. S4A) and then nascent DNA strands were separated through a sequencing polyacrylamide gel (Fig. S4B). Replication of the AP site plasmid generated nascent DNA strands at the −1 position (one nucleotide before the AP site; Fig. S4B, lanes 7-10) and these products were not detected during replication of control DNA (Fig. S4B, lanes 1-6). Importantly, there was no detection of −30 stall products (Fig. S4B, lanes 7-10) that could correspond to the footprint of the replicative helicase (58,64). Thus, the AP site stalls DNA polymerases, but not the replicative helicase, consistent with previous results (36).

### Leading and lagging strand AP sites stall DNA synthesis

In our initial experiments replication initiation was distributed across the plasmid template (64,74), replication forks encountered the AP site on either the leading or lagging strand template, and fork convergence happened rapidly. To control whether AP sites were encountered on the leading or lagging strand template and prevent fork convergence, AP sites were engineered into either DNA strand in a plasmid that contained an array of 50 *lacO* sequences. Addition of LacR led to the formation of a ‘LacR array’ that was previously shown to block fork movement and ensure that a lesion placed next to the array is encountered by a single fork from one direction (65). This resulted in plasmid templates where AP sites would be replicated on either the leading or lagging strand template (Fig. S5A,E).

We first replicated plasmid DNA containing a leading strand AP site. Replication intermediates were purified and then restriction-digested before being separated on a denaturing agarose gel. This allowed us to examine nascent strands from the fork that encountered the AP site (Fig. 2A). Replication of control DNA resulted in a transient stall product (Fig. 2B lane 1), as previously observed (Fig. 1F lanes 1-2). In contrast, replication of the leading strand AP resulted in a prominent stall intermediate (Fig. 2A, Fig. 2B lanes 6-8, Fig. 2C, Fig. S5D), which declined over approximately 30 minutes (Fig. 2B, lanes 7-8). Loss of the stall intermediate corresponded to the appearance of full-length strands that arose when replication forks bypassed the damage and encountered the LacR array (Fig. 2A, Fig. 2B lanes 6-10). Importantly, the generation of essentially all full-length nascent strands was impaired during replication of a leading strand AP site compared to control DNA (Fig. 2C, Fig. S5D) indicating that the AP site interfered with completion of all nascent DNA strands. Delayed completion of nascent strands was specific to the AP site because overall rates of DNA synthesis (Fig. S5B-C) and arrival of opposing replication forks at the other side of the LacR array (Fig. 2B) were unaffected compared to control DNA. These data show that the leading strand AP site causes persistent replication fork stalling that is ultimately bypassed, resulting in replication restart.

**Figure 2:**
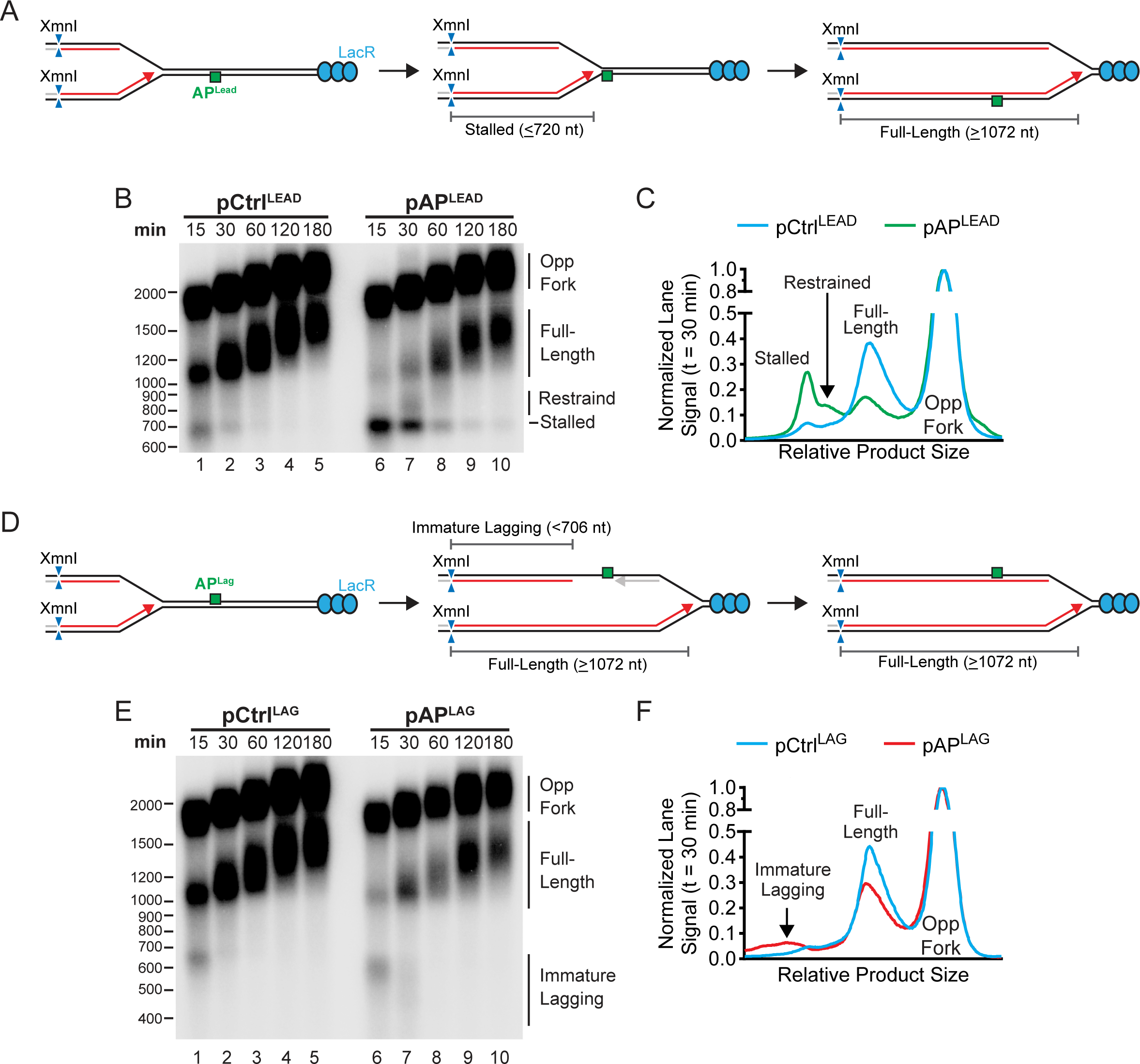
Leading and lagging strand AP sites stall DNA synthesis. A) Plasmid DNA harboring a leading strand AP site and LacR array was replicated in *Xenopus* egg extracts alongside control DNA. [α-32P]-dATP was added to label newly-synthesized DNA strands. Purified replication intermediates were then digested with XmnI to analyze nascent DNA strands. Products generated by the opposing fork (Opp Fork) by this analysis are shown in Fig. S5A. B) Nascent DNA strands from (A) were separated on an alkaline denaturing agarose gel then visualized by autoradiography. C) Lane profiles from (B) at 30 minutes. Lane profiles were normalized to the peak signal of Opp Fork. An independent experimental replicate is shown in Fig. S5D. D) Plasmid DNA harboring a lagging strand AP site was replicated and processed as in (A). Products generated by the opposing fork (Opp Fork) by this analysis are shown in Fig. S5E. E) Nascent DNA strands from (D) were separated on an alkaline denaturing agarose gel then visualized by autoradiography. F) Lane profiles from (E) at 30 minutes. Lane signals were normalized to the peak signal of Opp Fork. An independent experimental replicate is shown in Fig. S5H.

We next replicated plasmid DNA containing a lagging strand AP site and analyzed replication intermediates (Fig. 2D). As with the leading strand AP site, a transient stall was initially present during replication of both the lagging strand AP site plasmid and control DNA due to the LacR flanking the AP site (Fig. 2E lanes 1,6). Replication of the lagging strand AP gave rise to immature lagging strands (Fig. 2D, Fig. 2E lane 7) that were missing from the control (Fig. 2E lane 2). These strands were heterogenous in size, as expected for lagging strand stall products due to stochastic variation between molecules in the location of Okazaki fragment priming (Fig. 2F, Fig. S5H). These immature lagging strands arose from lagging strand stalling due to the presence of the AP site because overall rates of DNA synthesis (Fig. S5F-G) and arrival of opposing replication forks at the other side of the LacR array (Fig. 2E, Fig. S5E) were unaffected compared to control DNA.

### Leading strand AP sites rapidly stall replication of both leading and lagging strands

Replication of a leading strand AP site stalled most nascent strands (Fig. 2B, lanes 6-7), suggesting that both leading and lagging strand synthesis was stalled. To investigate this in more detail, we separately examined progression of leading and lagging strands during replication of a leading strand AP site.

Analysis of leading strands revealed a prominent stall intermediate which declined over approximately 30 minutes (Fig. 3A, Fig. 3B lanes 6-8, Fig. 3C, Fig. S6A). Replication of control DNA led to the expected transient leading strand stall (Fig. 3B lane 1, Fig. 3C, Fig. S6A). The transient stall products for control DNA disappeared by 30 minutes but most stall products for a leading strand AP site were still present at 30 minutes and did not resolve until later (Fig. 3C, Fig. S6A). This showed that essentially all leading strands were stalled by the leading strand AP site. By 120 minutes, most stalled leading products were resolved (Fig. 3C, Fig. S6A) and full-length strands reached a similar level to control DNA (Fig. 3D, Fig. S6B), demonstrating that most leading strands were able to bypass the leading strand lesion albeit with an approximately 45-minute delay. Thus, a leading strand AP site stalls most leading strands and restart mechanisms can efficiently bypass the AP site.

**Figure 3:**
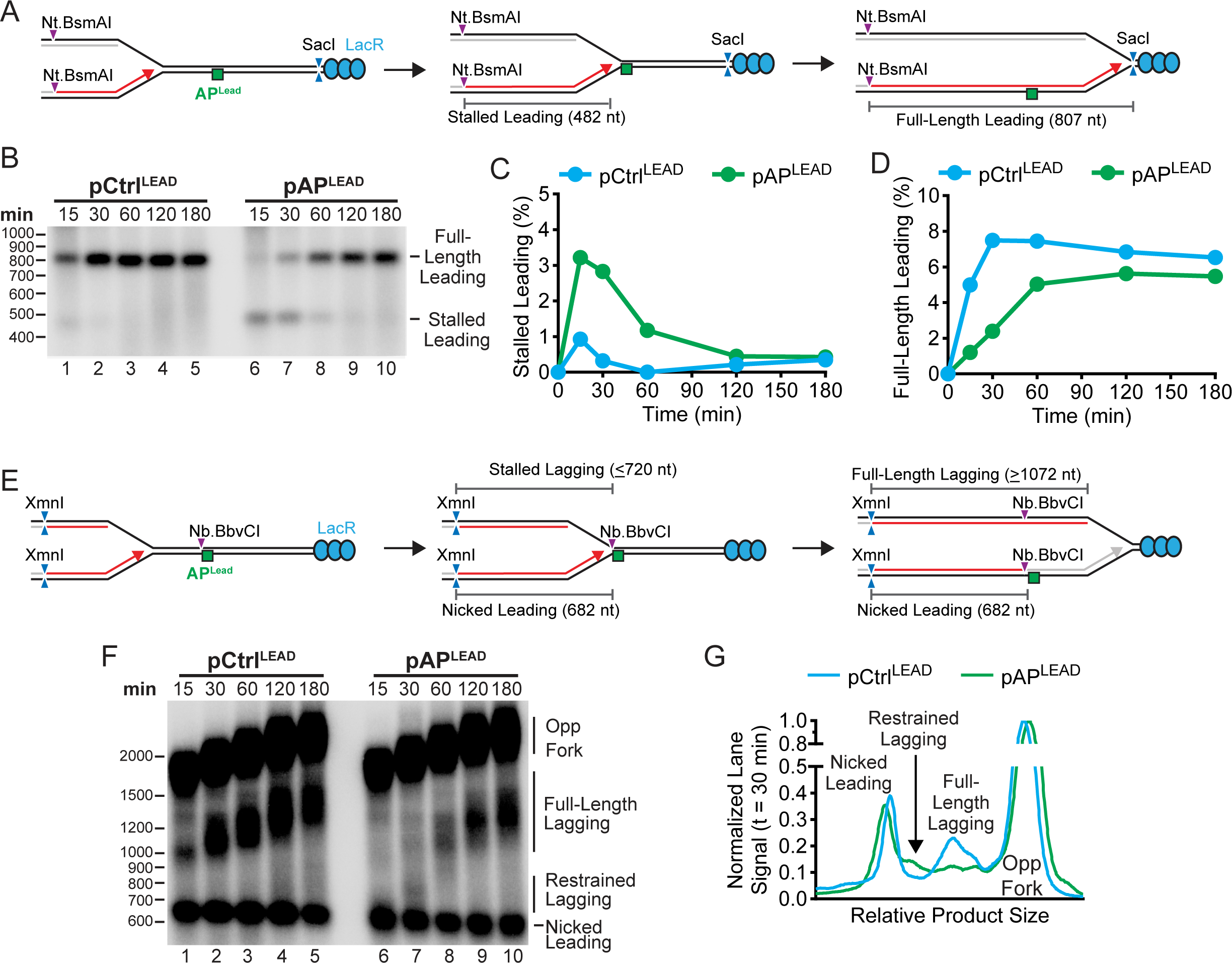
Leading strand AP sites stall both nascent DNA strands. A) Plasmid DNA harboring a leading strand AP site and LacR array was replicated in *Xenopus* egg extracts alongside control DNA. [α-32P]-dATP was added to label newly-synthesized DNA strands. Purified replication intermediates were then digested with Nt.BsmAI and SacI to analyze nascent leading strands. B) Nascent leading strands from (A) were separated on an alkaline denaturing agarose gel then visualized by autoradiography. C) Quantification of stalled leading strands from (B). An independent experimental replicate is shown in Fig. S6A. D) Quantification of full-length leading strands from (B). An independent experimental replicate is shown in Fig. S6B. E) Purified replication intermediates from (A) were digested with XmnI and Nb.BbvCI to analyze nascent lagging strands. F) Nascent strands from (E) were separated on an alkaline denaturing agarose gel then visualized by autoradiography. G) Lane profiles from (F) at 30 minutes. Lane signals were normalized to the peak signal of Opp Fork. An independent experimental replicate is shown in Fig. S6C.

Analysis of lagging strands arising from the leading strand AP site revealed two distinct populations (Fig. 3E-F). At 30 minutes, most lagging strands were ∼100 nucleotides longer than expected if stalled opposite the leading strand AP site (Fig. 3F-G, Fig. S6C-D), but shorter than the full-length product that arose from completely synthesized lagging strands that encountered the LacR array (Fig. 3F-G, Fig. S6C). These products corresponded to lagging strands that were extended beyond the AP site but had stalled or slowed (Fig. S6D). Because leading strands were largely stalled at the AP site during this time (Fig. 3C, Fig. S6A) this indicated that limited uncoupling of nascent DNA strand synthesis had taken place (i.e. lagging strands were extended only ∼100 nucleotides beyond the AP site, while leading strands remained stalled at the AP). Lagging strands eventually restarted and reached full-length by 60 minutes (Fig. 3E-F), correlating with restart of leading strands (Fig. 3C-D). Thus, at a leading strand AP site, replisomes quickly stall and then restart at the same time that leading strands bypass the lesion.

Replisome stalling was limited when forks converged at an AP site (Fig 1A-D), in contrast to the pronounced replisome stalling observed at a leading strand AP site (Fig. 3E-G). This was notable because approximately 50% of AP sites should be replicated on the leading strand template if fork convergence is not blocked by a LacR array (58,64). Thus, our data show that a converging fork can rescue leading strand stalling. Lagging strands can be extended past the AP site following a leading strand stall (Fig. 3E-G) and gapped molecules are ultimately generated (Fig 1A-B), which indicates that the converging causes termination downstream of the AP site followed by gap filling once the AP site is bypassed.

### Lagging strand AP sites stall nascent lagging but not leading nascent strands

Nascent DNA strands were minimally inhibited by a lagging strand AP site (Fig. 2D-F), suggesting that efficient bypass of leading or lagging strands, or both, took place. To investigate this in more detail, we separately examined progression of leading and lagging strands during replication of a lagging strand AP site.

Analysis of lagging strands revealed a prominent lagging strand stall intermediate which declined over approximately 30 minutes (Fig. 4A, Fig. 4B lanes 6-8, Fig. 4C). A transient leading strand stall was also visible at the 15-minute time point for both the lagging strand AP site plasmid and control DNA (Fig. 4A, Fig. 4B lane 1 and 6). Importantly, lagging strand stall products on the AP site plasmids were stable for approximately 30 minutes and then resolved by 60 minutes (Fig. 4C). This matched the timing of formation and resolution of leading strand stall products at a leading strand AP site (Fig. 3C, Fig. S6A). Thus, bypass of the AP site on the damaged strand occurred at a similar speed regardless of whether the AP site was on the leading or lagging strand.

Analysis of leading strands arising from a lagging strand AP site (Fig. 4D) did not detect any stall products (Fig. 4E). In fact, extension of leading strands beyond the lagging strand AP was indistinguishable from control DNA (Fig. 4E-F, Fig. S5E). Thus, a lagging strand AP site does not appear to affect synthesis of leading DNA strands. Because only leading strand AP sites stall synthesis on the opposite undamaged strand (Fig. 3F-G, Fig. S6C, Fig. 4E-F, Fig. S6E), these data show that leading and lagging strand AP sites exert different effects on replication fork progression beyond the lesion.

### Bypass of AP sites requires TLS

Nascent strands stalled one nucleotide away from the AP site immediately prior to bypass (Fig. S4), which is consistent with TLS (36). To test whether TLS synthesis is required for bypass of an AP site we sought to inhibit REV1, which acts as both a TLS polymerase and scaffold for other TLS polymerases (75,76). To this end, we examined replication of a leading strand AP site plasmid in extracts that were mock-immunodepleted or REV1-immunodepleted (Fig 5A, Fig. S7A). In mock-depleted extracts, stalled leading strand products accumulated at the AP site and then declined at the same time as full-length leading strands appeared (Fig. 5B lanes 11-15, Fig. 5C-D), as previously observed (Fig. 3A-D). In REV1-depleted extracts, stalled leading strand products accumulated to the same extent, but did not decline, and there was greatly reduced accumulation of full-length leading strand products (Fig. 5B, lanes 16-20, Fig. 5C-D,). Although the quantification approach underestimated the abundance of stall products due to nucleotide content, no appreciable difference was observed when a correction for this was applied. (Fig. S7D-E). The lack of full-length leading strand products reflected a specific defect in bypass of the AP site because REV1-depleted extracts exhibited unaltered total DNA synthesis (Fig. S7B-C). Furthermore, there was no difference in formation of full-length products when control DNA was replicated in mock-or REV1-depleted extracts (Fig. 5B lanes 1-10, Fig. 5D). Overall, immunodepletion of REV1 blocks bypass of the AP site, suggesting that TLS is crucial for AP site bypass.

**Figure 4:**
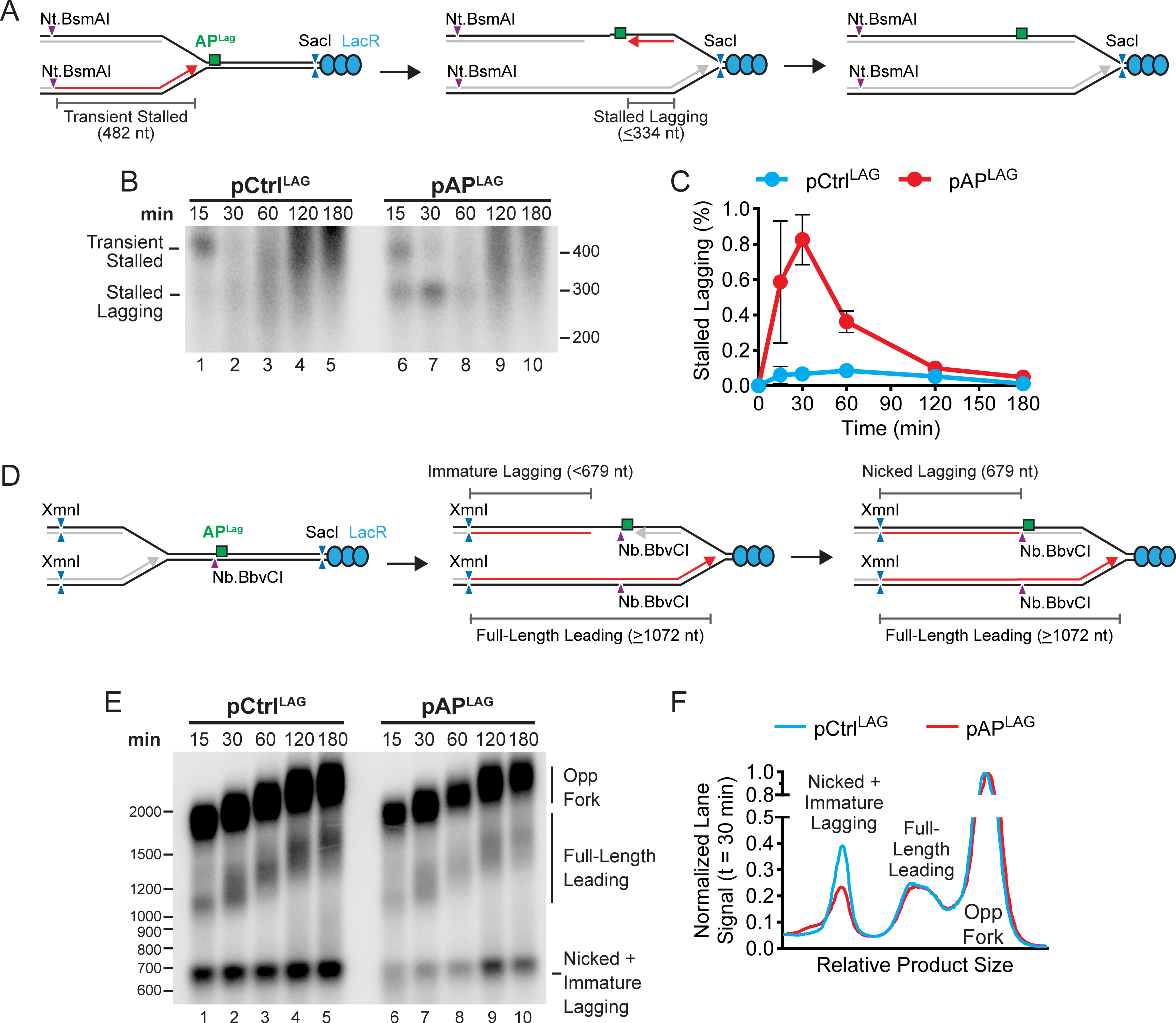
Lagging strand AP sites only stall nascent lagging strands. A) Plasmid DNA harboring a lagging strand AP site and LacR array was replicated in *Xenopus* egg extracts alongside control DNA. [α-32P]-dATP was added to label newly-synthesized DNA strands. Purified replication intermediates were then digested with Nt.BsmAI and SacI to analyze nascent lagging strands. Transient leading strands stalling is expected to arise from the LacR flanking the AP site and the expected leading strand products are indicated. B) Nascent strands from (A) were separated on an alkaline denaturing agarose gel then visualized by autoradiography. C) Quantification of stalled lagging strands from (B). Mean ± s.d., *n* = 3 independent experiments. D) Purified replication intermediates from (A) were digested with XmnI and Nb.BbvCI to analyze nascent leading strands. E) Nascent strands from (E) were separated on an alkaline denaturing agarose gel then visualized by autoradiography. F) Lane profiles from (E) at 30 minutes. Lane signals were normalized to the peak signal of Opp Fork. An independent experimental replicate is shown in Fig. S6E.

**Figure 5:**
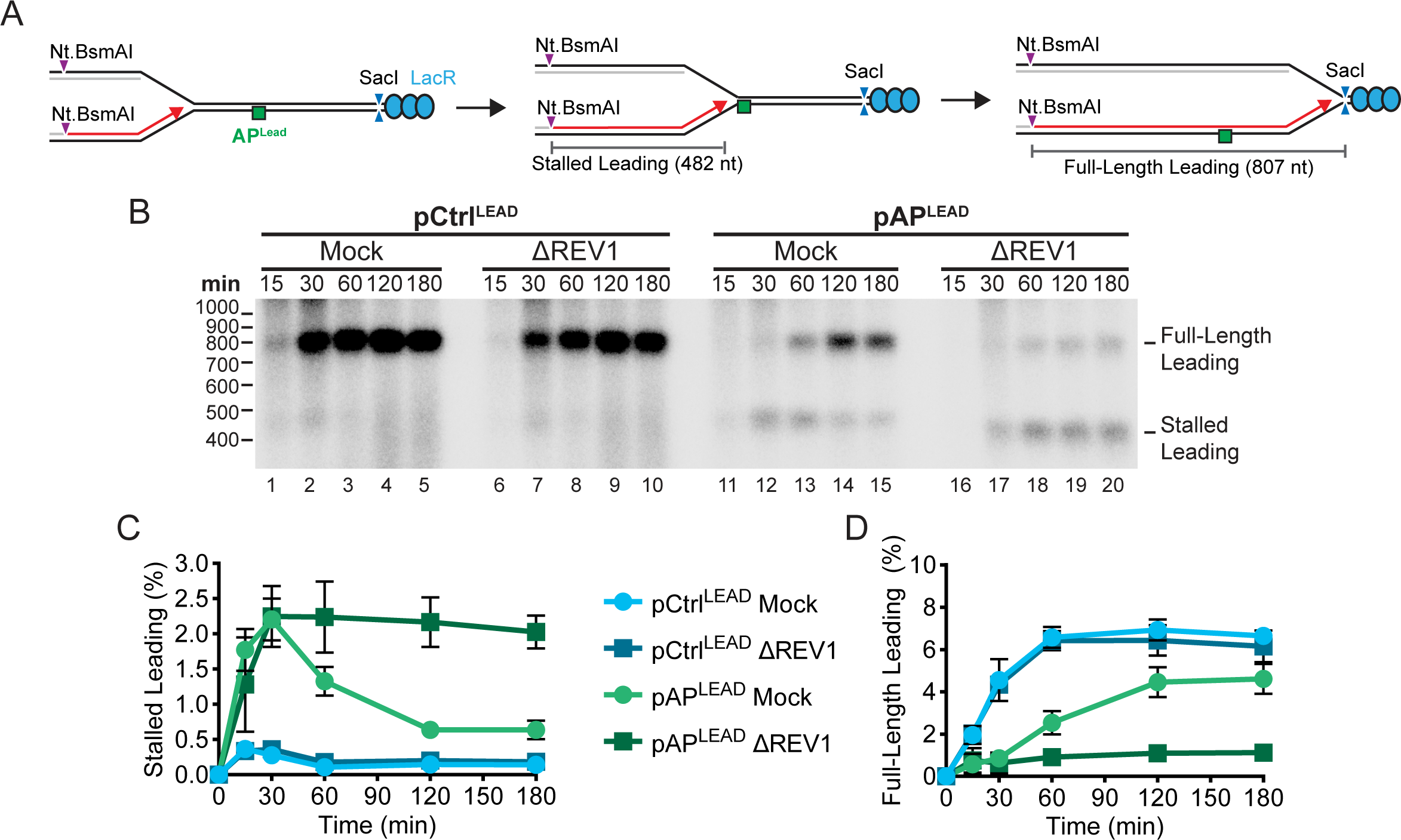
Immunodepletion of Rev1 impairs bypass of a leading strand AP site. A) A leading strand AP site was replicated in Mock-or REV1-immunodepleted *Xenopus* egg extracts alongside control DNA. [α-32P]-dATP was added to label newly-synthesized DNA strands. Purified replication intermediates were then digested with Nt.BsmAI and SacI to analyze nascent leading strands. B) Nascent strands from (A) were separated on an alkaline denaturing agarose gel then visualized by autoradiography. C) Quantification of stalled leading strands from (B). Mean ± s.d., *n* = 3 independent experiments. See Fig. S7D for quantification of stalled leading strands as a percentage of total nascent leading strands. D) Quantification of full-length leading strands from (B). Mean ± s.d., *n* = 3 independent experiments. See Fig. S7E for quantification of full-length leading strands as a percentage of total nascent leading strands.

Because it was not feasible to rescue the REV1-immunodepletion using purified recombinant protein due to the size of REV1 and its numerous binding partners (75–77), we performed additional interventions to further test whether TLS is required for AP site bypass. We first immunodepleted the TLS DNA polymerase POLκ (Fig. S8A), which is required for TLS in *Xenopus* egg extracts (78). POLκ-depletion had no effect on overall DNA replication (Fig. S8B-D) but inhibited resolution of stalled products and limited formation of full-length products (Fig. S9). We next inhibited ubiquitin signaling, which is also required for TLS (79). To this end, extracts were treated with the de-ubiquitinating enzyme inhibitor ubiquitin vinyl sulfone (Ub-VS) to deplete ubiquitin, as previously described (64,80). Treatment with Ub-VS did not affect overall DNA synthesis (Fig. S10A-C) but did block resolution of stall products and formation of full-length products (Fig. S10D-I). Considering that POLκ-depletion, inhibition of ubiquitin signaling, and REV1-depletion impaired bypass, we conclude that TLS is crucial for bypass of a leading template strand AP site.

To test whether TLS is also required for bypass of lagging template strand AP sites, we analyzed the requirement for REV1 when a lagging strand AP site was replicated. REV1-depletion did not affect overall DNA synthesis (Fig. S11A-C) but did block resolution of stalled lagging strands (Fig. S10D-G). The same result was observed when POLκ was depleted (Fig. S12). Overall, these data indicate that the mechanism used to bypass the AP site is the same regardless of whether the lesion is on the leading or lagging template strand.

### Leading strand AP bypass supports lagging strand restart

Our observations that lagging strands were temporarily stalled by a leading template strand AP site (Fig. 3E-G) and lagging strand restart correlated with restart of leading strands (Fig. 3A-D) suggested that bypass of the leading template strand AP site might be important to facilitate lagging strand restart. To test this possibility, we examined lagging strand progression beyond a leading template strand AP site under conditions where AP site bypass was blocked by depletion of REV1 (Fig. 6A). Replication of control DNA was identical for mock-or REV1-depleted extracts (Fig. 6B lanes 1-10, Fig. 6C, Fig. S13A), as expected. Replication of the leading template strand AP plasmid in either mock-or REV1-depleted extracts initially produced lagging strands ∼100 nucleotides beyond the AP site (Fig. 6B, lanes 13,18), as previously observed (Fig. 3E-F, Fig. S6D). In mock-depleted extracts, lagging strands were ultimately extended to full-length strands (Fig. 6B lanes 14-15). In contrast, approximately 50% of lagging strand products progressed to no more than ∼200 bp downstream of the AP site and persisted for up to 120 minutes in REV1-depleted extracts (Fig. 6B, lanes 19-20, Fig. 6D, Fig. S13B). Similar results were obtained by depleting POLκ, which also caused a reduction in the formation of full-length lagging strands and persistence of shorter, incomplete products for at least 120 minutes (Fig. S13C-G). Thus, TLS-dependent bypass of the leading strand AP site is important to restart and extend replication of the undamaged lagging strands, which otherwise remain stalled for several hours.

**Figure 6:**
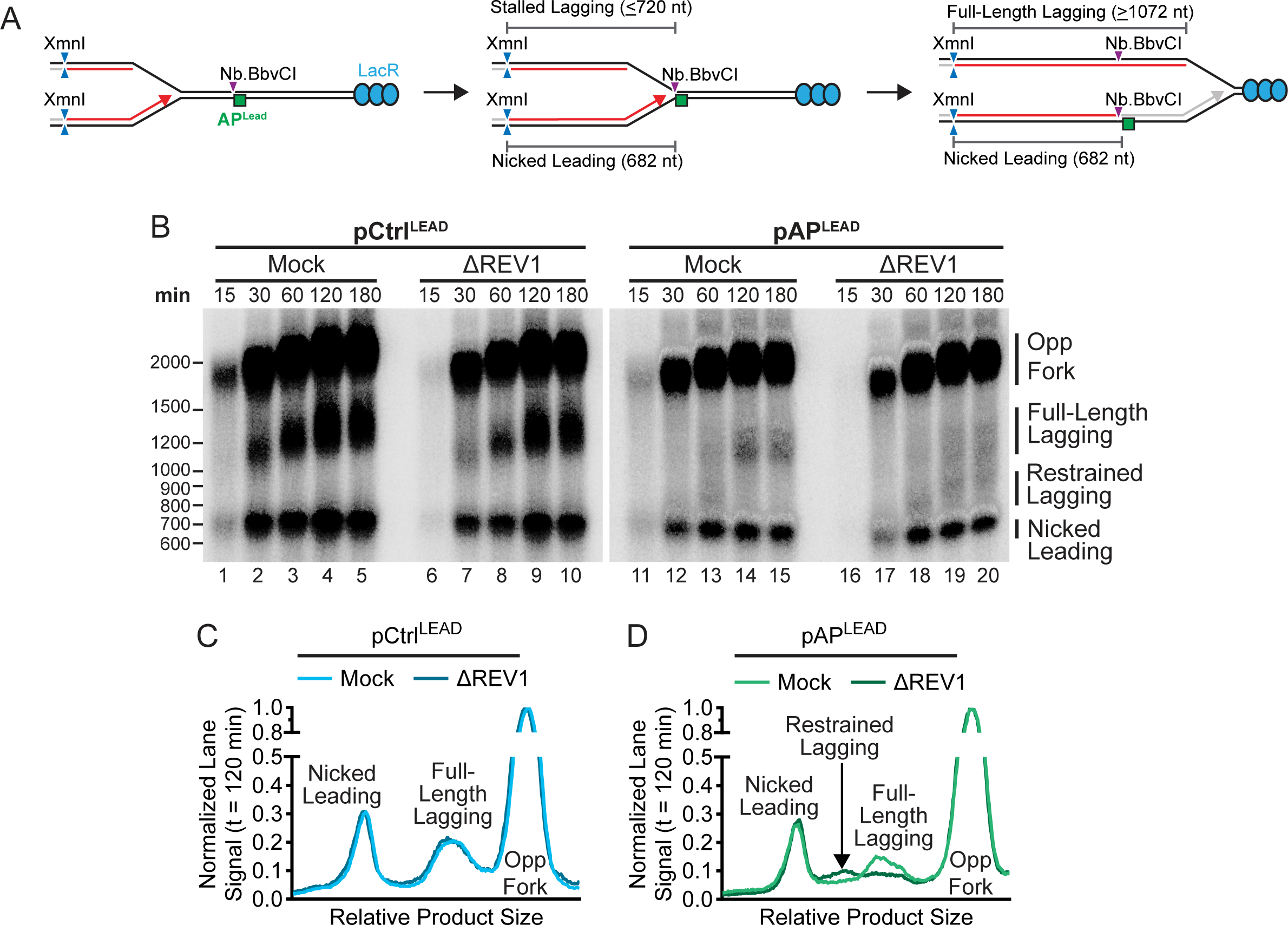
Impaired bypass of a leading strand AP site restrains lagging strand synthesis. A) A leading strand AP site was replicated in Mock-or REV1-immunodepleted *Xenopus* egg extracts alongside control DNA. [α-32P]-dATP was added to label newly synthesized DNA strands. Purified replication intermediates were then digested with Nb.BbvCI and XmnI to analyze nascent lagging strands. B) Nascent strands from (A) were separated on an alkaline denaturing agarose gel then visualized by autoradiography. Images shown are different exposures of the same gel because overall signal was lower for pAP^LEAD^. C) Lane profiles from (B) for control DNA at 30 minutes. Lane signals were normalized to the peak signal of Opp Fork. Lane signals were normalized to Ctrl. An independent experimental replicate is shown in Fig. S13A. D) Lane profiles from (B) for AP-containing DNA at 30 minutes. Lane signals were normalized to the peak signal of Opp Fork. An independent experimental replicate is shown in Fig. S13B.

## Discussion

Using an approach to study replication of site-specific AP sites we have generated data that supports a detailed model for how AP sites affect replication forks (Fig. 7). Encounter with a leading strand AP initially causes replisomes to stall ∼100 bp downstream of the lesion (Fig. 7A). Upon lesion bypass by TLS, replisome progression resumes (Fig. 7A). In contrast, encounter with a lagging strand AP site does not result in detectable replisome slowing or stalling (Fig. 7B). Instead, Okazaki fragment repriming generates a post-replicative gap that is filled in following TLS across the lesion (Fig. 7B).Therefore, lagging strand AP site bypass happens at a post-replicative gap, while leading strand bypass happens proximal to the replication fork, consistent with on-the-fly TLS (52). Importantly, leading and lagging strand AP site bypass occur with similar timing and involve the same proteins, consistent with a recent report of shared mechanisms for bypass of base modification on the leading and lagging strand template (81). Thus, leading and lagging strand AP sites elicit analogous bypass mechanisms despite exerting different effects on replisome progression. Importantly, our data establishes that these mechanisms (Fig. 7) apply during replication of AP sites themselves rather than their derivatives (SSBs, DPCs, ICLs) (12–16).

**Figure 7:**
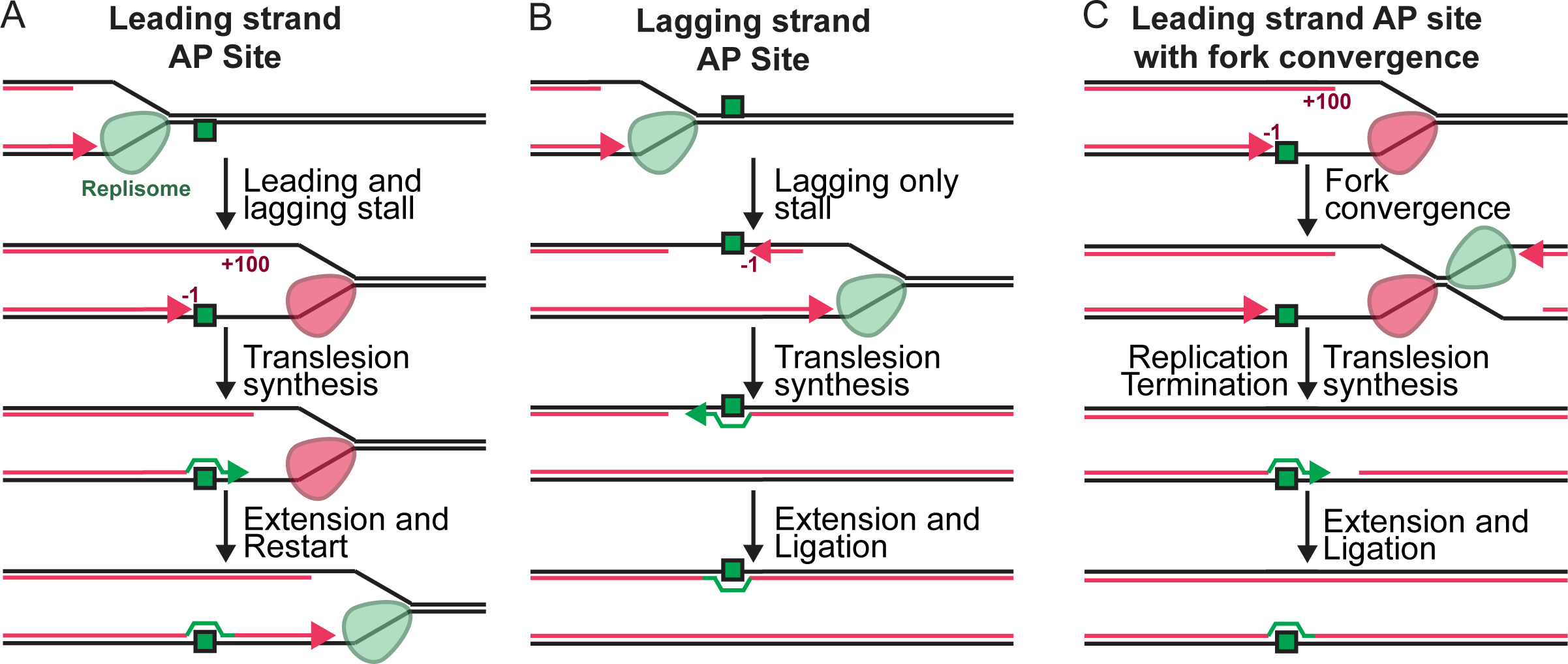
Model for AP site replication in *Xenopus* egg extracts. A) Encounter with a leading strand AP site causes nascent leading strands to stall one nucleotide before the lesion (−1). Nascent lagging strands are restrained and stall approximately 100 nucleotides downstream of the lesion, resulting in limited uncoupling. The AP site is then bypassed by TLS, which results in restart of both leading and lagging strand synthesis. B) Encounter with a lagging strand AP site causes nascent lagging strands to stall one nucleotide before the lesion. Nascent leading strands are unaffected and continue synthesis. The AP site is then bypassed by TLS, which results in fill-in of the post-replicative gap. C) Fork convergence allows for a leading strand AP to by bypassed without restart of leading and lagging strand synthesis. If a converging fork arrives prior to restart, then replication termination takes place immediately downstream of the AP site. Bypass of the AP site is followed by extension of the leading strand to the downstream lagging strand of the opposing fork, to which it is then ligated.

Our results also show that fork convergence at an AP site results in fully separated daughter molecules that contain a post-replicative gap (Fig. 1). Prior to fork convergence, the AP site will be initially encountered on either the leading or lagging strand template. Lagging strand AP sites are bypassed to leave a post-replicative gap (Fig. 7B), and fully separated daughter molecules will arise from termination downstream of the AP site. Leading strand AP sites cause fork stalling (Fig. 7A), which can be rescued by a converging fork (Fig. 7C). In this case, termination will take place immediately downstream of the AP site, which will result in formation of a post-replicative gap and separation of daughter DNA molecules (Fig. 7C). Importantly this scheme is different to replication of strand-specific DPCs where forks converge at the lesion and transiently stall, irrespective of which strand is damaged, presumably due to the bulky nature of the DPC (58). Thus, leading and lagging strand AP sites can both result in a post-replicative gap, but through different mechanisms and these events differ from those that take place at bulkier DNA adducts.

Our observations of how AP sites affect vertebrate replication align with the effects of non-bulky DNA damage on purified yeast replisomes (35,61), but also reveal unexpected differences. Broadly, both yeast and vertebrate replisomes are unaffected by lagging strand lesions which are readily tolerated by repriming downstream of the damage (35,61). Vertebrate replisomes stall ∼100 bp downstream of the lesion in *Xenopus* egg extracts. In contrast, purified yeast replisomes did not undergo detectable slowing within ∼650 nucleotides of the damage (35). Uncoupled replisomes are expected to slow because leading strand synthesis is required for optimal helicase unwinding (32–34). However, these experiments predict only ∼5-10-fold slowing of replisome progression (32–34). In contrast, we find that when bypass of the leading strand AP is blocked, lagging strands remain stalled for several hours. Thus, our observations indicate that replisome activity is much more dramatically inhibited than previously thought. In the future it will be important to address whether rapid stalling reflects an intrinsic property of the replisome or other components of the DNA damage response, such as ATR signaling (82–85).

In our experiments, AP site bypass was blocked by depletion of REV1 or POLκ, regardless of which strand contained the lesion site. The reliance on TLS for bypass is consistent with previous studies in *Xenopus* egg extracts that reported that TLS is the major pathway for bypass of DPCs (86), ICLs (67), as well as major and minor groove lesions (78). It is also consistent with a recent report that non-bulky DNA damage in human cells is bypassed by TLS on both strands (81). Nonetheless, this observation is surprising because multiple DNA polymerases are capable of AP site bypass (37–44,47,48), and template switching should also be able to promote bypass (30,31,87). It is possible that all TLS is heavily dependent on the scaffolding activity of REV1 and/or POLκ, as suggested for other DNA lesions (78). Furthermore, although *Xenopus* egg extracts support both fork reversal (62,88) and homologous recombination (89,90), which could allow template switching to take place, it may be infrequent or suppressed. Template switching is best characterized downstream of a DNA double-strand break (91). We did not observe double-strand breaks (Fig. 1) or template switching (Fig. 5) so it is possible that template switching is rare without double strand break formation. In the absence of TLS, replisomes stalled at a leading strand AP should be able to restart by repriming (30,31,61,92) or other TLS-independent mechanisms (11,45,46). However, we did not observe TLS-independent restart (Fig. 6), and it will be important to determine why this was the case. Despite its potential for generating mutations, TLS may be advantageous because it does not risk large-scale copy number variations and other chromosomal alterations that are attributable to template switching (30,31).

In conclusion, we have developed an approach to examine strand-specific replication of DNA templates containing stabilized AP sites. This approach may be generally applicable to other lesions that can be converted to different types of DNA damage. Unexpectedly, we found rapid replisome stalling upon encounter with a leading strand AP site, and restart largely depended on TLS bypass of the lesion. Our studies provide an avenue for future research to understand the underlying mechanisms that control replisome movement and choice of replication stress tolerance mechanism.

## Data Availability

All data is available within the manuscript.

## Supplementary Data Statement

Supplementary Data are available at NAR Online

## Acknowledgements

We thank Johannes Walter for REV1 antibodies.

## Author contributions

D.C. and J.M.D. conceived of and supervised the project. M.C. completed all experiments. S.D. assisted with development of experimental approaches and data interpretation. The manuscript was written by J.M.D and M.C. with editing by D.C.

---

**Figure S1:**
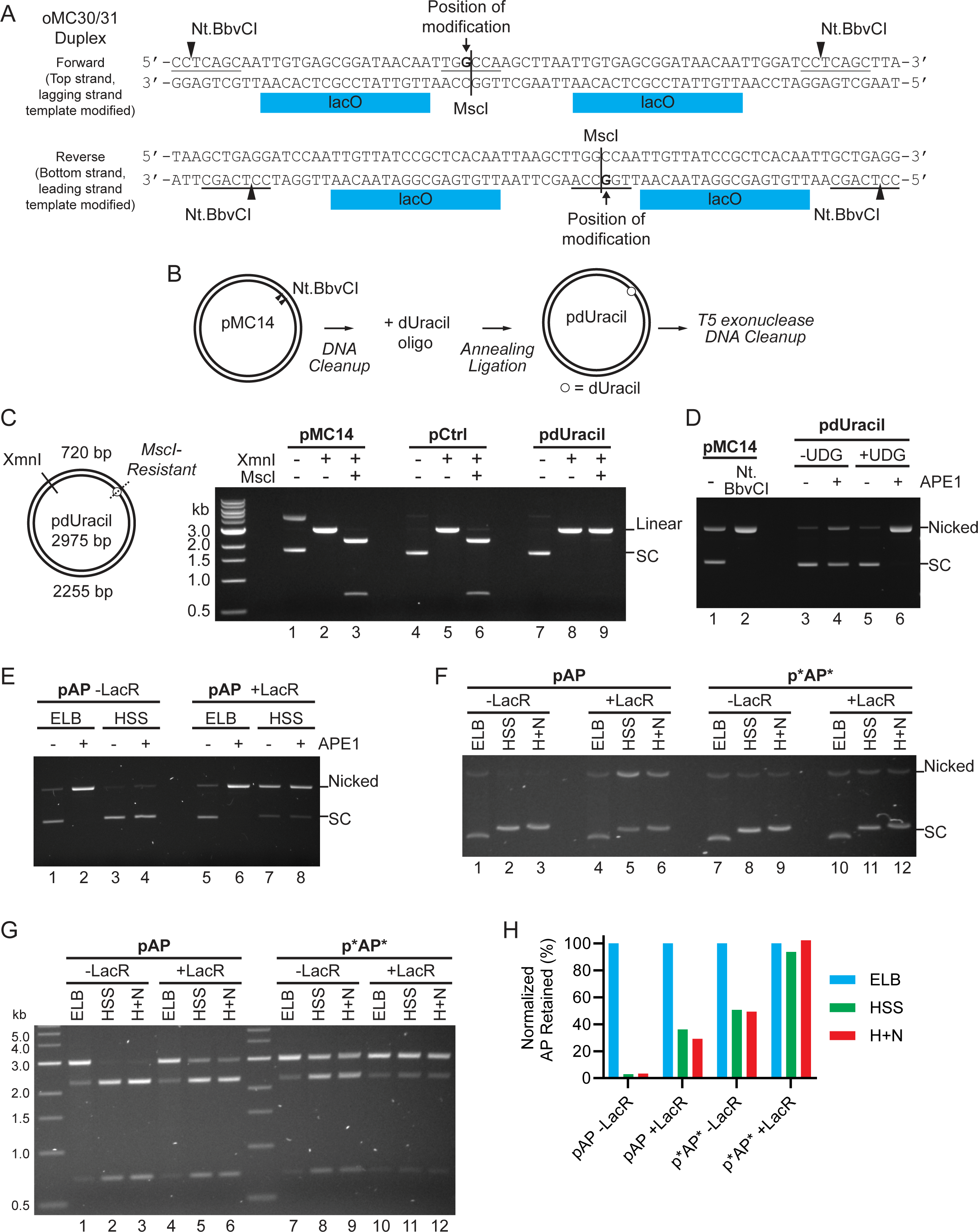
Construction and stability of AP site plasmids. A) DNA oligonucleotides oMC30 and oMC31 were annealed to form a blunt duplex which was cloned into the PsiI restriction site of pJD161. This insert contains Nt.BbvCI restriction sites placed 63 nucleotide (nt) apart on the same strand. Incorporation of the oMC30/31 duplex in the forward direction generated pMC9, which allows for for incorporation of a modified oligonucleotide between the Nt.BbvCI sites on the Top strand. Incorporation of the oMC30/31 duplex in the reverse direction generated pMC10 and pMC14, which allow for incorporation of a modified on the Bottom strand. Replication forks can approach the modified region of pMC14 from either direction and thus encounter the modified strand on either the leading or lagging strand template (Fig.1). pMC9 and MC10 harbor a *lacO* array downstream of the indicated sequence to ensure the modified strand is encountered on the lagging (pMC9) or leading (pMC10) strand template when incubated in LacR to assemble a replication barrier. B) To insert a site- and strand-specific AP site into plasmid DNA, pMC14 was first nicked with Nt.BbvCI. A DNA oligonucleotide containing a site-specific dUracil was then annealed and ligated into the gap between nicking sites. Excess unligated oligos and nicked plasmid intermediates were then removed and the remaining fully ligated plasmid was purified. C) pMC14 was modified with a Uracil-containing oligonucleotide (pdUracil) or an unmodified control oligo (pCtrl), as in (B), then digested with XmnI and MscI to screen for incorporation of the dUracil. XmnI linearizes the plasmid into a 3 kb fragment to eliminate differences between plasmids that arise from different levels of supercoiling. MscI targets the same position as the inserted dUracil modification, which blocks digestion by MscI. Resistance to digestion by MscI acts as a read out for incorporation of the site-specific dUracil modification. pdUracil was resistant to MscI (compare lane 9 with lanes 3 and 6), which confirmed that dUracil was efficiently incorporated. D) pdUracil was treated with UDG to generate plasmids containing an AP site (pAP),then treated with APE1 to test for the generation of an AP site. APE1 converts AP sites to single-stranded DNA breaks. Thus, conversion of supercoiled plasmids to nicked species indicates the presence of AP sites. Parental plasmid (pMC14) was included as a marker for supercoiled (SC) DNA, and also nicked with Nt.BbvCI as a marker for nicked plasmid. Treatment with UDG and APE1 resulted in predominantly nicked plasmids (lane 6), which demonstrated efficient conversion of dUracil to AP sites. E) To evaluate the stability of pAP in *Xenopus* egg extracts, plasmid DNA containing an abasic site (pAP) was incubated in High Speed Supernatant (HSS) or buffer control (Egg Lysis Buffer, ELB) then treated with APE1 to assess the presence of the AP site. pAP became resistant to APE1 following incubation with HSS (compare lanes 2 and 4), which indicated that the AP site was repaired by *Xenopus* egg extracts. To test whether LacR binding adjacent to the AP site could confer stability in *Xenopus* egg extracts, pAP was pre-incubated with LacR before being incubated in HSS. Most plasmids became nicked under these conditions (compare lanes 5 and 7), which indicated that the LacR-bound AP site was incised in HSS but not fully repaired. Thus, LacR-binding conferred partial stability to the AP site in *Xenopus* egg extracts. F) Phosphorothioate-modified AP site plasmid (p*AP*) was incubated in *Xenopus* egg extracts with or without LacR to evaluate stability of the AP site compared to pAP, which lacked phosphorothioates. In addition to High Speed Supernatant (HSS) of *Xenopus* egg extracts, HSS plus NucleoPlasmic Extract (NPE)) combined (H+N) was also used to mimic the final extract concentration during a typical replication reaction using *Xenopus* egg extracts. pAP incubated with LacR became nicked (compare lanes 5-6 with lane 4), p*AP* did not (compare lanes 11-12 with lane 10). Thus, p*AP* incubated with LacR was stabilized in *Xenopus* egg extracts compared to pAP. G) To evaluate whether p*AP* retained the AP site in *Xenopus* egg extracts, Products from (F) were purified then digested with XmnI and MscI (as in (C)). The ∼3000 bp and ∼700 bp fragments of XmnI-MscI digestion (C) increased when pAP was incubated with LacR (compare lanes 5-6 with lane 4), but not when p*AP* was incubated with LacR (compare lanes 11-12 with lane 10). Note that a background level of MscI digest products is detected even incubation with *Xenopus* egg extracts (lanes 1, 4, 7, 10) because the AP site was not fully resistant to MscI, in contrast to dUracil (in (C)). Note also that presence of the AP site could not be evaluated by treatment with APE1 (as in (D)-(E)) because phosphorothioate modifications confer resistance to APE1. Overall, these data indicate that the combination of phosphorothioates and LacR binding stabilize AP sites in *Xenopus* egg extracts. H) To quantitatively assess AP site retention in *Xenopus* egg extracts, retention of the ∼3000 bp product from (G) was quantified. Values were normalized to the ELB condition to account for partial sensitivity of the AP site to MscI. The combination of phosphorothioates and LacR (p*AP*+LacR) resulted in no appreciable loss of AP sites. This demonstrated that AP sites could be stabilized in *Xenopus* egg extracts.

**Figure S2:**
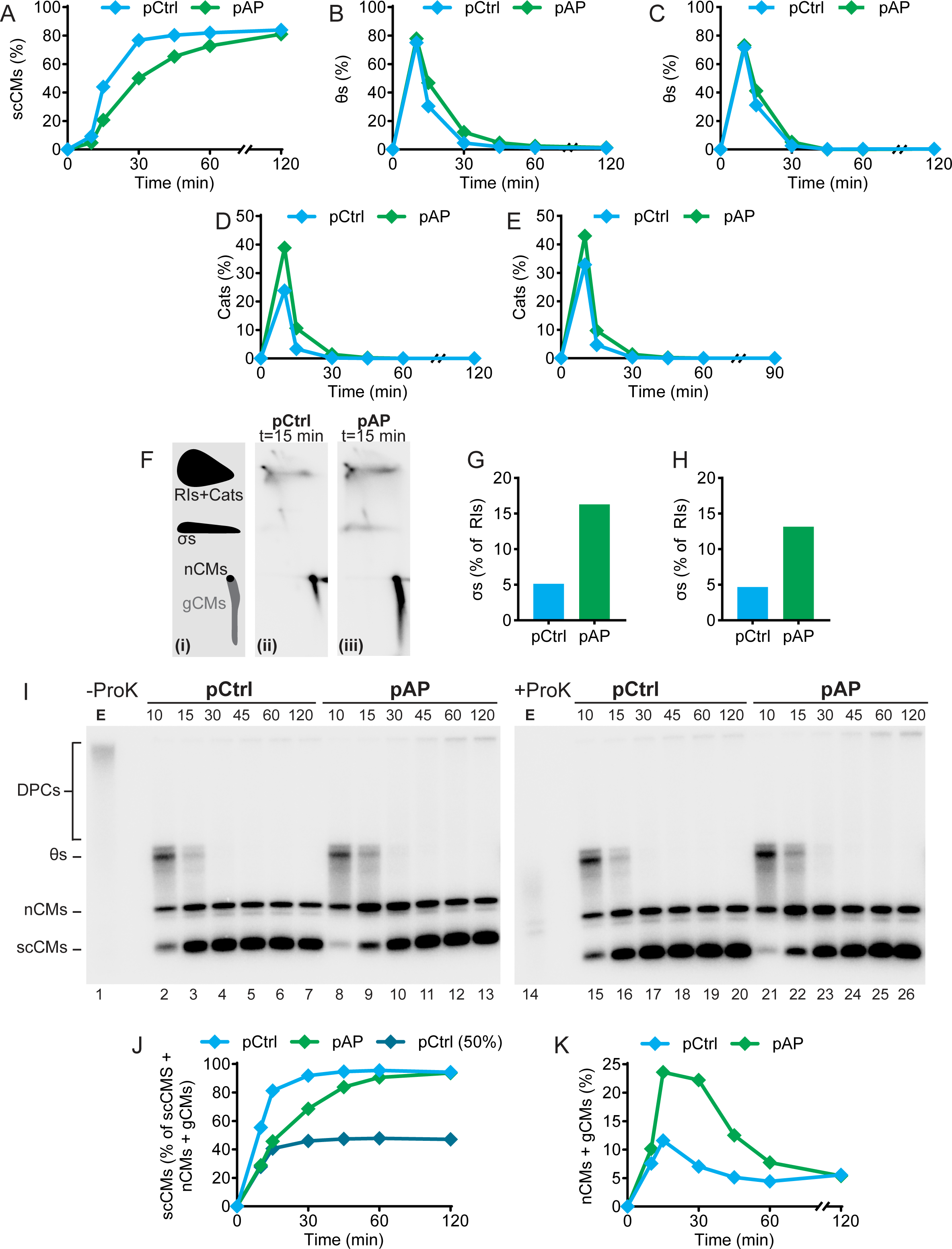
Analysis of products generated during replication of control and AP site plasmids. A) Independent experimental replicate of Fig. 1C. B) Quantification of thetas (θs) from Fig. 1B. C) Independent experimental replicate of (B) D) Quantification of catenanes (cats) from Fig. 1B. E) Independent experimental replicate of (D) F) Products from Fig. 1A at 15 minutes were separated by 2D gel electrophoresis to visualize sigmas (σs), replication intermediates and catenanes (RIs+Cats), nicked Circular Monomers (nCMs), and gapped Circular Monomers (gCMs). G) Quantification of σs from (F). H) Independent experimental replicate of (G). I) To assess the presence of DNA-protein cross-links (DPCs), replication products were generated as in Fig. 1B, treated with or without Proteinase K, then separated on a native agarose gel and visualized by autoradiography (as in (58)). Replication of control DNA in the presence of etoposide (100 μM, E) was used as a positive control for DPC formation. Etoposide treatment resulted in a noticeable decrease in mobility when Proteinase K treatment was omitted (compare lanes 1 and 14) due to DPC formation, as expected (93). In contrast, omission of Proteinase K had little effect on DNA structures arising from replication of pCTRL or pAP (compare lanes 2-13 and 15-26). Thus, there was no evidence of DPC formation during replication of pAP. J) Independent experimental replicate of Fig. 1D. K) Quantification of supercoiled circular monomers (scCMs) as a percentage of total circular monomers (including nicked and gapped circular monomers (nCMs and gCMs)) from Fig. 1B to estimate frequency of stalling at AP sites. Replication of a strand-specific DNA lesion in *Xenopus* egg extracts is expected to result in one fully replicated daughter molecule (scCM) and one nicked or gapped daughter molecule (nCM or gCM) (58). Thus, if all AP sites are retained and stall DNA synthesis, half the daughter molecules are expected to be replicated with the same timing as undamaged plasmid DNA (pCtrl). To estimate timing of replication for a single undamaged DNA strand, half of the signal from the control condition was plotted (pCtrl (50%)). At 10- and 15-minutes signal from replication of pAP closely matched signal from pCtrl (50%), consistent with half of DNA strands being replicated with normal timing in the pAP condition. The datapoints then diverged once pCtrl (50%) plateaued, consistent with half of DNA strands being replicated much more slowly in the pAP condition. Thus, these data suggest that most AP sites are retained and stall replication on one strand only.

**Figure S3:**
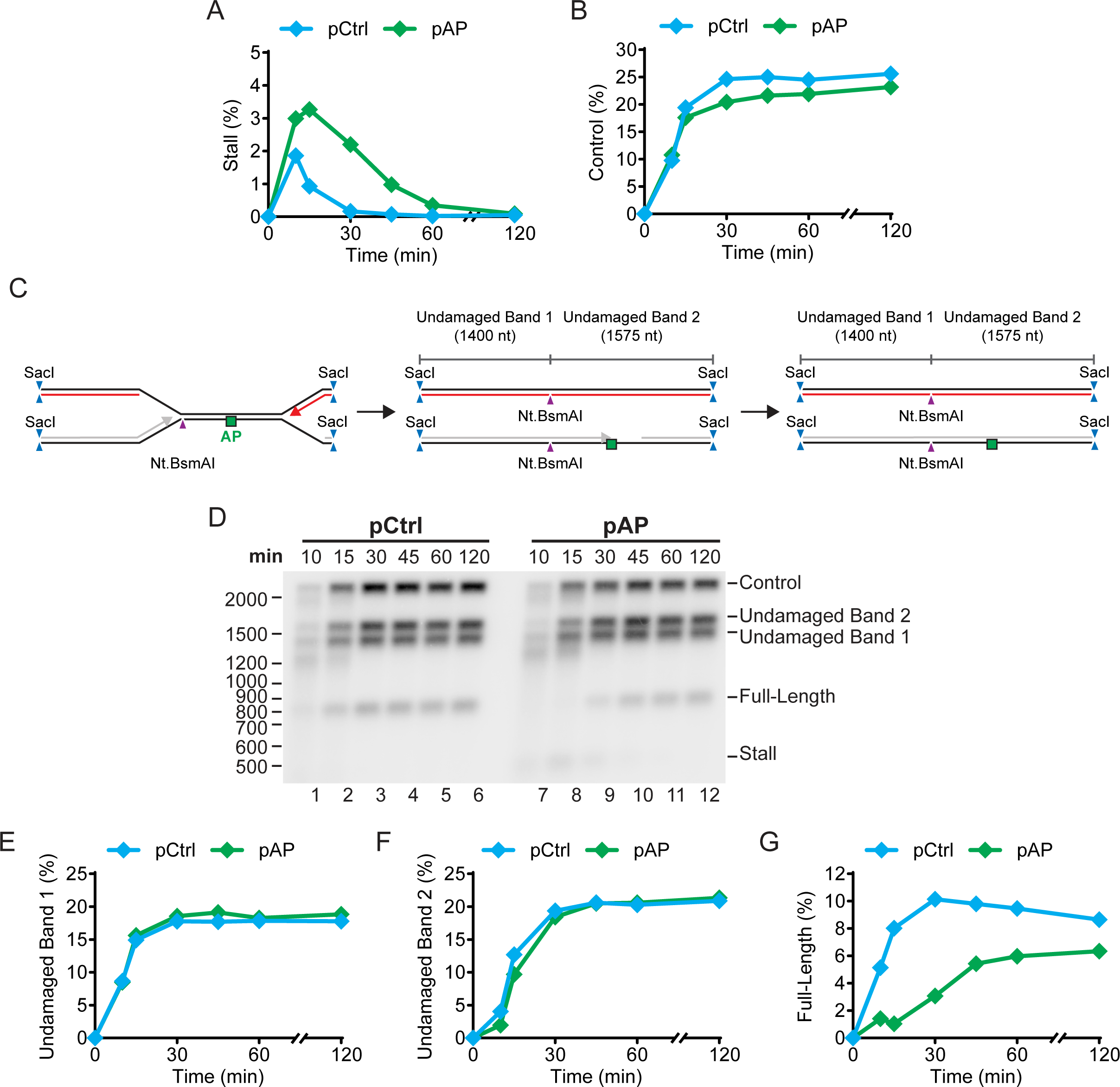
Analysis of nascent strands generated during replication of control and AP site plasmids. A) Independent experimental replicate of Fig. 1G B) Quantification of Control products from Fig. 1F. C) Purified replication intermediates from Fig. 1A were digested with Nt.BsmAI and SacI as in Fig. 1E. Digestion products arising from the undamaged strand (not shown in Fig. 1E) are shown. D) Expanded gel image of Fig. 1F showing products from replication of the undamaged strand. E) Quantification of Undamaged Band 1 from (D) F) Quantification of Undamaged Band 2 from (D) G) Independent experimental replicate of Fig. 1H.

**Figure S4:**
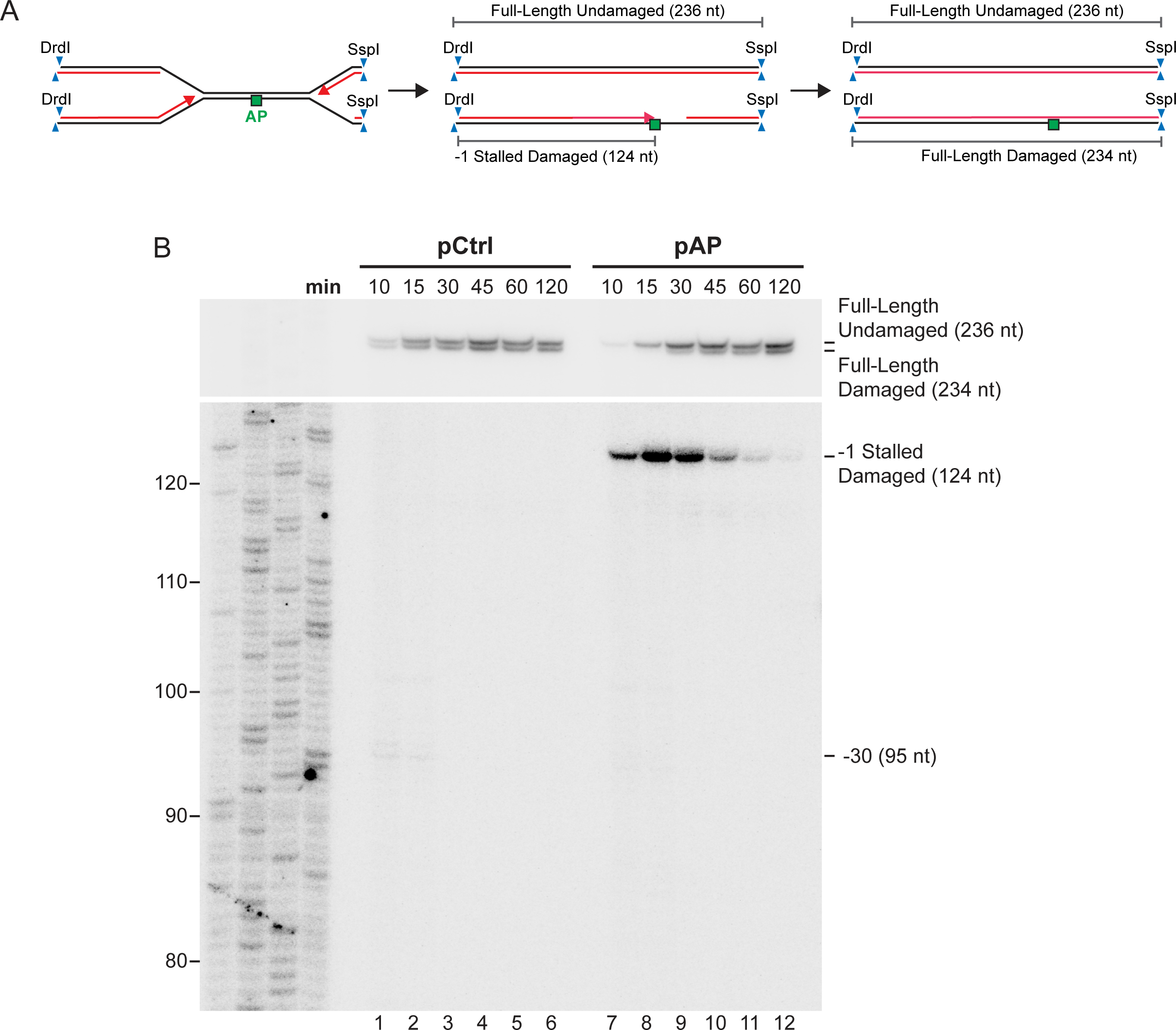
Abasic sites result in nascent strands stalled one nucleotide before the lesion. A) Purified replication intermediates from Fig. 1A were digested with DrdI and SspI to analyze stalled intermediates during replication of an abasic site. B) Nascent DNA strands from (A) were separated on a 6% sequencing polyacrylamide gel then visualized by autoradiography. Replisome stalling at the AP site would be expected to stall nascent leading strands approximately 30 nucleotides prior to the AP site (58,64), but no persistent stall products at this position (−30) were detected.

**Figure S5:**
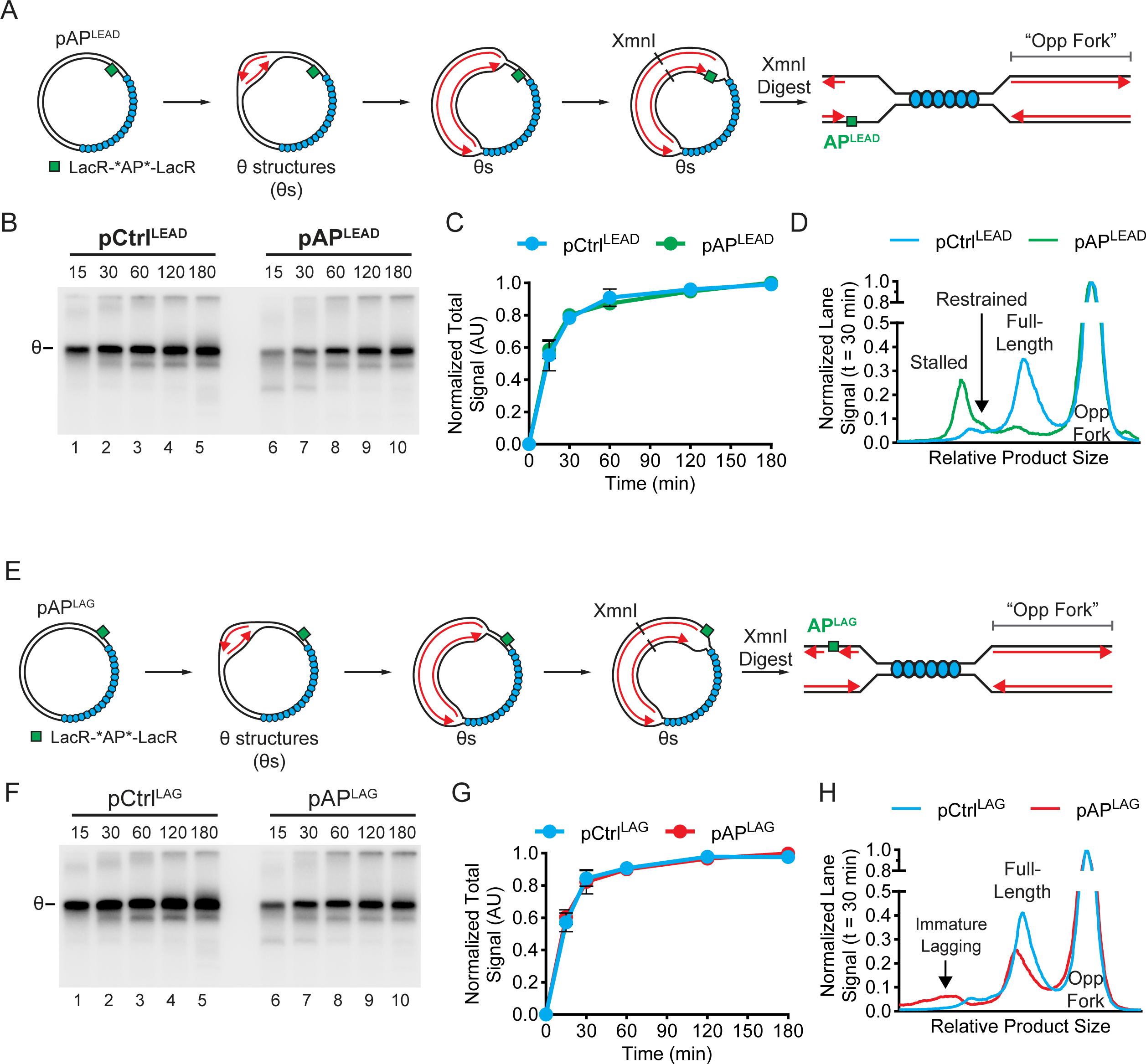
Replication of leading strand-specific and lagging strand-specific AP site plasmids. A) Alternate view of Fig. 2A that depicts the DNA structures formed during plasmid replication as well as the opposing fork (Opp Fork) that does not encounter the AP site and serves as a control. B) Undigested products from Fig. 2A were separated on a native agarose gel and visualized by autoradiography. C) Quantification total synthesis from (B). Values are normalized to the peak signal recorded for each template. Mean ± s.d., *n* = 3 independent experiments is shown. D) Independent experimental replicate of Fig. 2C. E) Alternate view of Fig. 2D that depicts the DNA structures formed during plasmid replication as well as the opposing fork (Opp Fork) that does not encounter the AP site and serves as a control. F) Undigested products from Fig. 2D were separated on a native agarose gel and visualized by autoradiography. G) Quantification total synthesis from (F). Values are normalized to the peak signal recorded for each template. Mean ± s.d., *n* = 3 independent experiments is shown. H) Independent experimental replicate of Fig. 2F.

**Figure S6:**
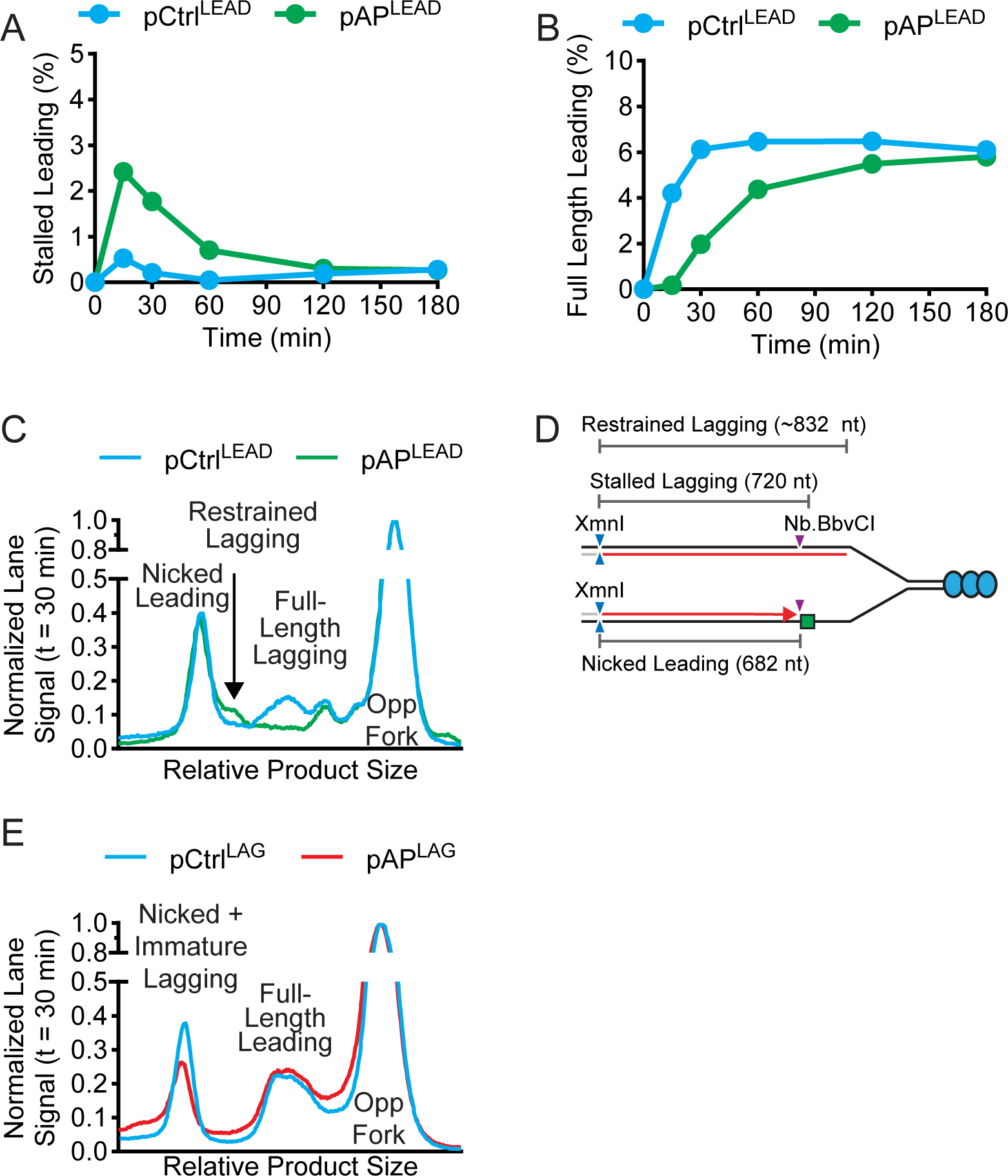
Replicate analyses of products formed during replication of leading and lagging strand AP sites. A) Independent experimental replicate of Fig. 3C. B) Independent experimental replicate of Fig. 3D. C) Independent experimental replicate of Fig. 3G. D) Schematic of restrained lagging strand intermediates formed during replication of leading strand AP sites (see Fig. 3E-G). Lagging strands were initially ∼150 nucleotides longer than the nicked leading strand products (Fig. 3F, lane 7). This indicated that the lagging strand products were approximately 832 nucleotides in size. Lagging strands stalled immediately prior to the AP site would be 720 nucleotides. Thus, the lagging strands were stalled ∼100 nucleotides beyond the lesion. E) Independent experimental replicate of Fig. 4F.

**Figure S7:**
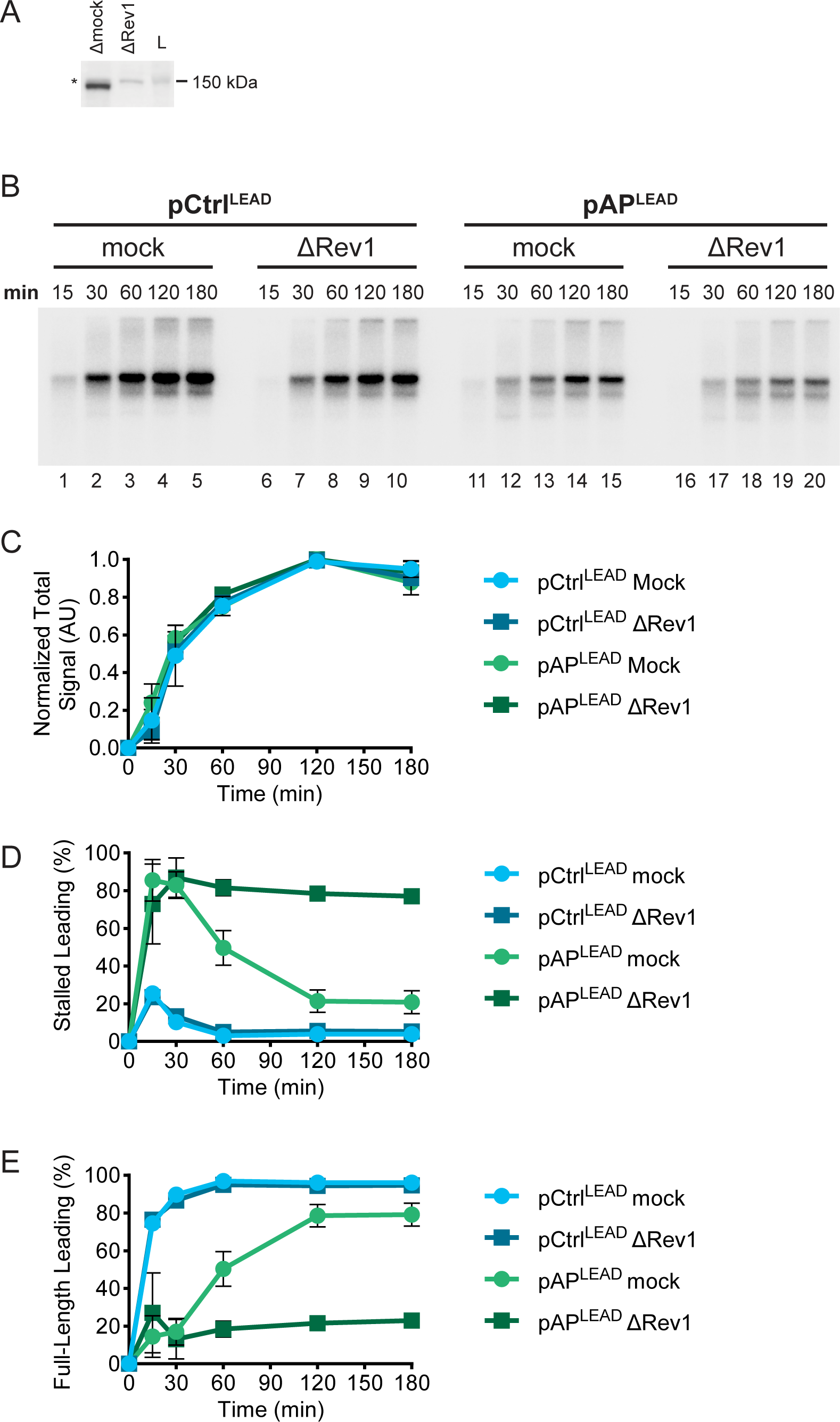
Immunodepletion of Rev1 has no effect on overall rates of DNA synthesis. A) Western blot of mock- and Rev1-depleted NPE. B) Undigested products from Fig. 5B were separated on a native agarose gel and visualized by autoradiography. C) Quantification total synthesis from (B). Values are normalized to the peak signal recorded for each condition. Mean ± s.d., *n* = 3 independent experiments is shown. D) Quantification of stalled leading strands from Fig. 5B as a percentage of total nascent leading strands. These values are corrected for expected incorporation of radiolabeled dATP according to dA content of the stalled and full-length products. Mean ± s.d., *n* = 3 independent experiments. E) Quantification of full-length leading strands from Fig. 5B as a percentage of total nascent leading strands. These values are corrected for expected incorporation of radiolabeled dATP according to dA content of the stalled and full-length products. Mean ± s.d., *n* = 3 independent experiments.

**Figure S8:**
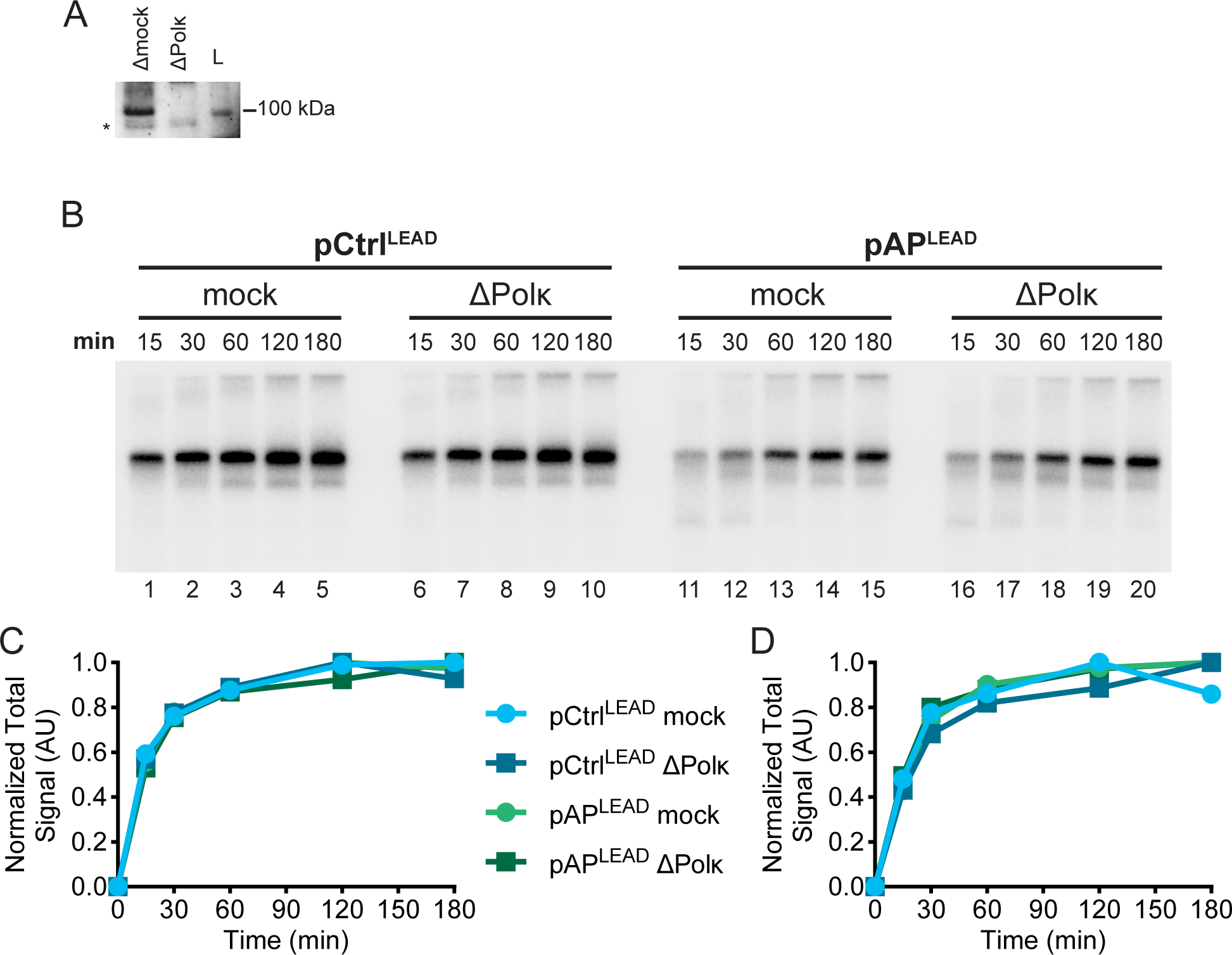
Immunodepletion of Polκ has no effect on overall rates of DNA synthesis. A) Western blot of mock- and Polκ-depleted NPE. B) Plasmid DNA was replicated as in Fig. 5A but using Polκ-depleted extracts. Undigested products were separated on a native agarose gel and visualized by autoradiography. C) Quantification total synthesis from (B). Values are normalized to the peak signal recorded for each condition. Mean ± s.d., *n* = 3 independent experiments is shown. D) Independent experimental replicate of (C).

**Figure S9:**
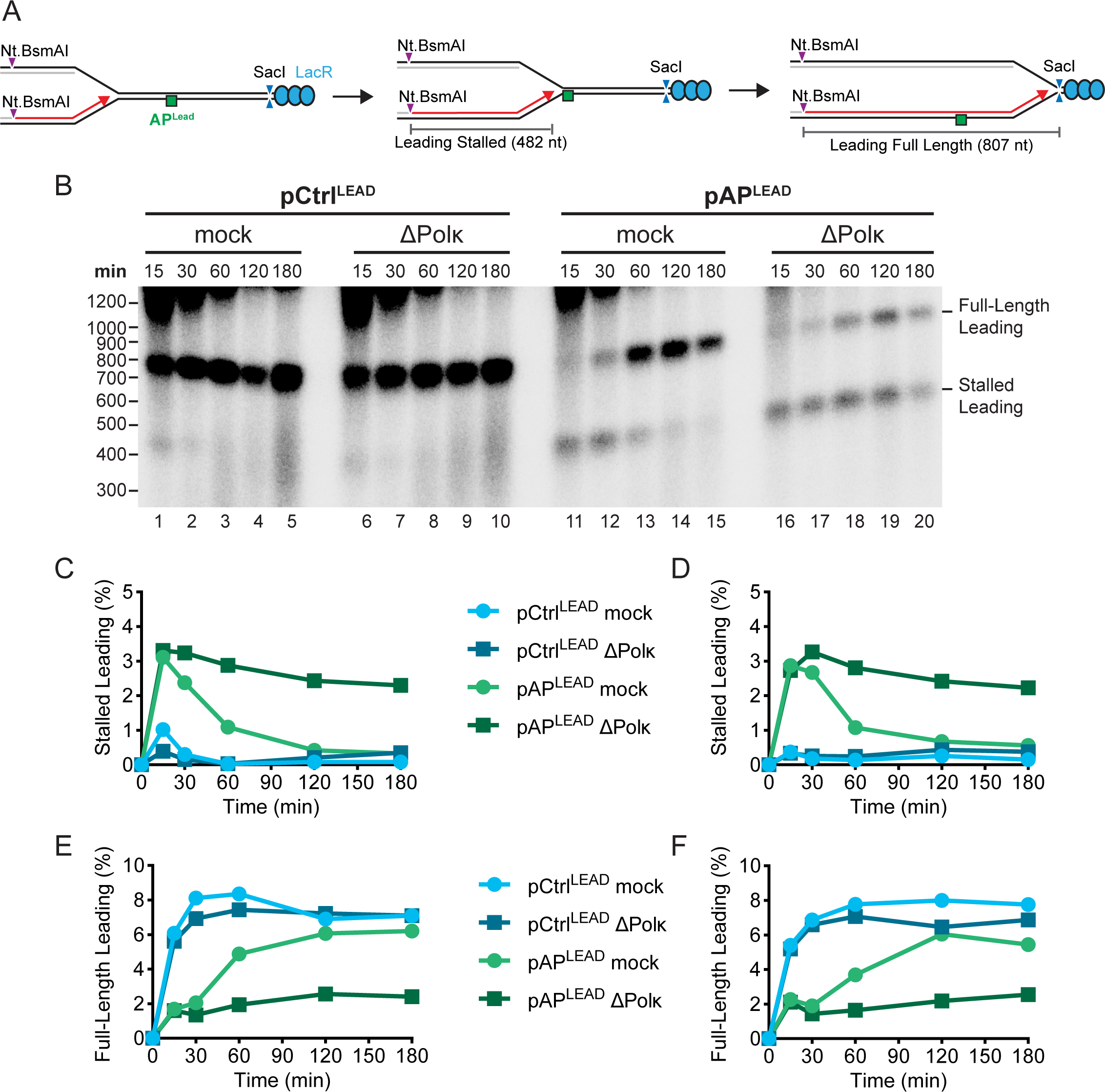
Immunodepletion of Polκ largely inhibits bypass of leading strand AP sites. A) Plasmid DNA was replicated as in Fig. 5A but using Polκ-depleted extracts. B) Nascent strands from (A) were separated on an alkaline denaturing agarose gel and visualized by autoradiography. C) Quantification of stalled leading strands from (B). D) Independent experimental replicate of (C). E) Quantification of full-length leading strands from (B). F) Independent experimental replicate of (E).

**Figure S10:**
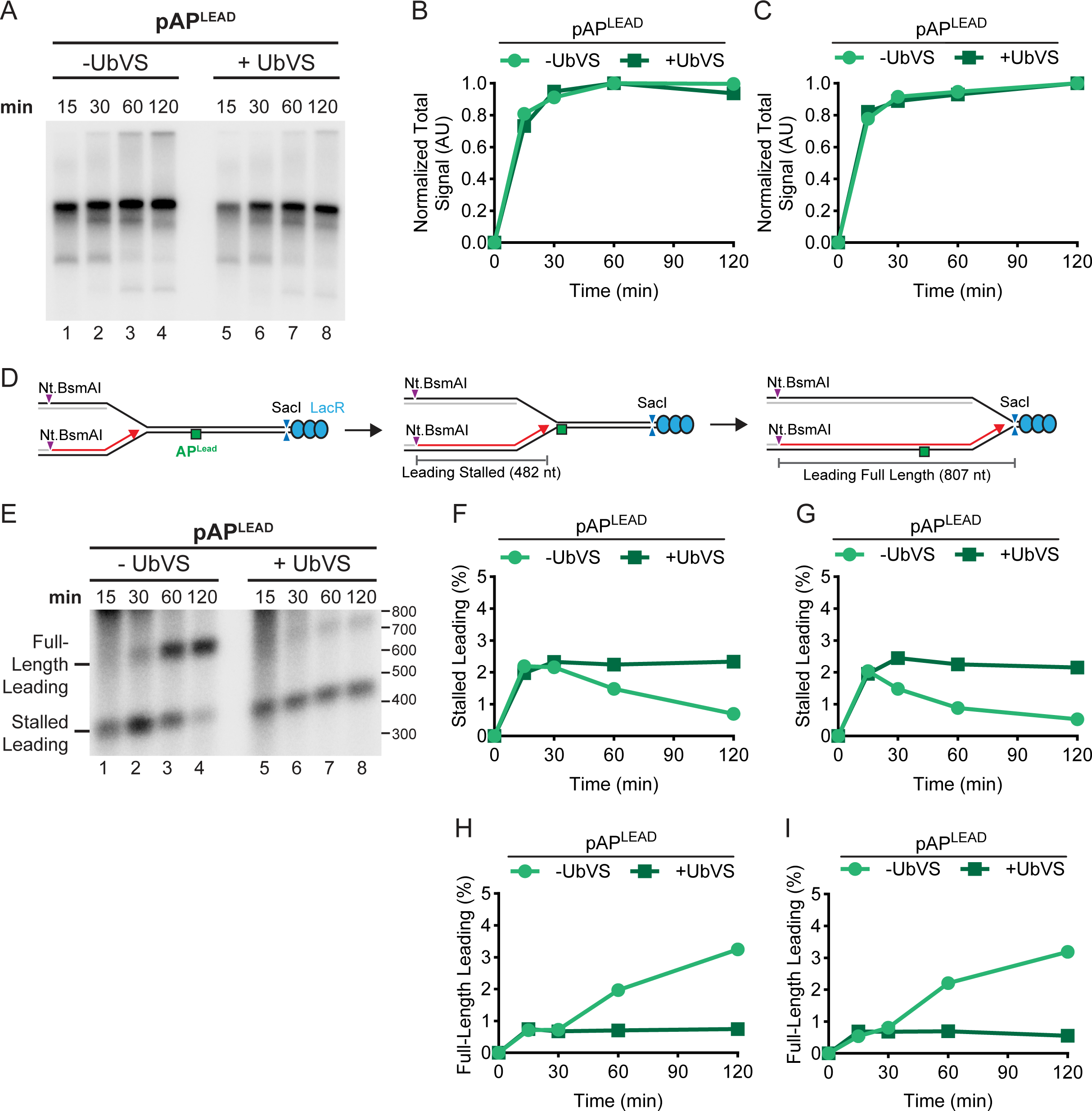
Ubiquitin depletion inhibits bypass of leading strand AP sites. A) Plasmid DNA harboring a leading strand AP site and LacR array was replicated in *Xenopus* egg extracts in the absence or presence of Ubiquitin Vinyl Sulfone (UbVS, 20 μM), which depletes free ubiquitin from *Xenopus* egg extracts(80). [α-32P]-dATP was added to label newly synthesized DNA strands. Products were resolved on a native agarose gel and visualized by autoradiography. B) Quantification total synthesis from (A). Values are normalized to the peak signal recorded for each condition. C) Independent experimental replicate of (B). D) Purified products from (A) were digested with Nt.BsmAI and SacI to detect nascent leading strands. E) Nascent strands from (D) were separated on an alkaline denaturing agarose gel and visualized by autoradiography. F) Quantification of stalled leading strands from (E). G) Independent experimental replicate of (F). H) Quantification of full-length leading strands from (E). I) Independent experimental replicate of (H).

**Figure S11:**
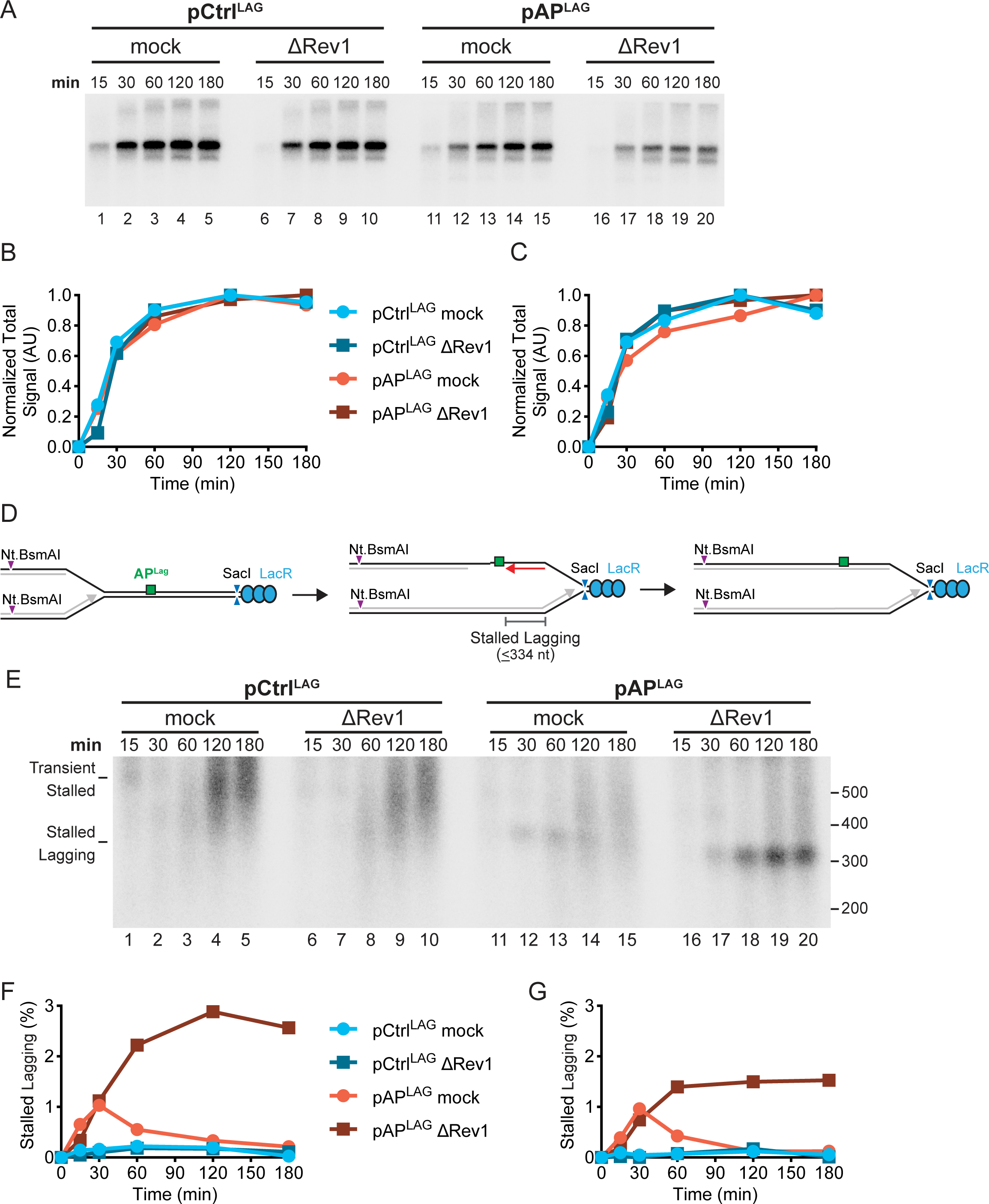
Immunodepletion of Rev1 impairs bypass of lagging strand AP sites. A) Plasmid DNA was replicated as in Fig 4A using mock- and REV1-depleted extracts. Undigested products were separated on a native agarose gel and visualized by autoradiography. B) Quantification total synthesis from (A). Values are normalized to the peak signal recorded for each condition. C) Independent experimental replicate of (B). D) Purified products from (A) were digested with Nt.BsmAI and SacI to detect nascent lagging strands. E) Nascent strands from (D) were resolved on an alkaline denaturing agarose gel and visualized by autoradiography. F) Quantification of stalled lagging strands from (E). G) Independent experimental replicate of (F).

**Figure S12:**
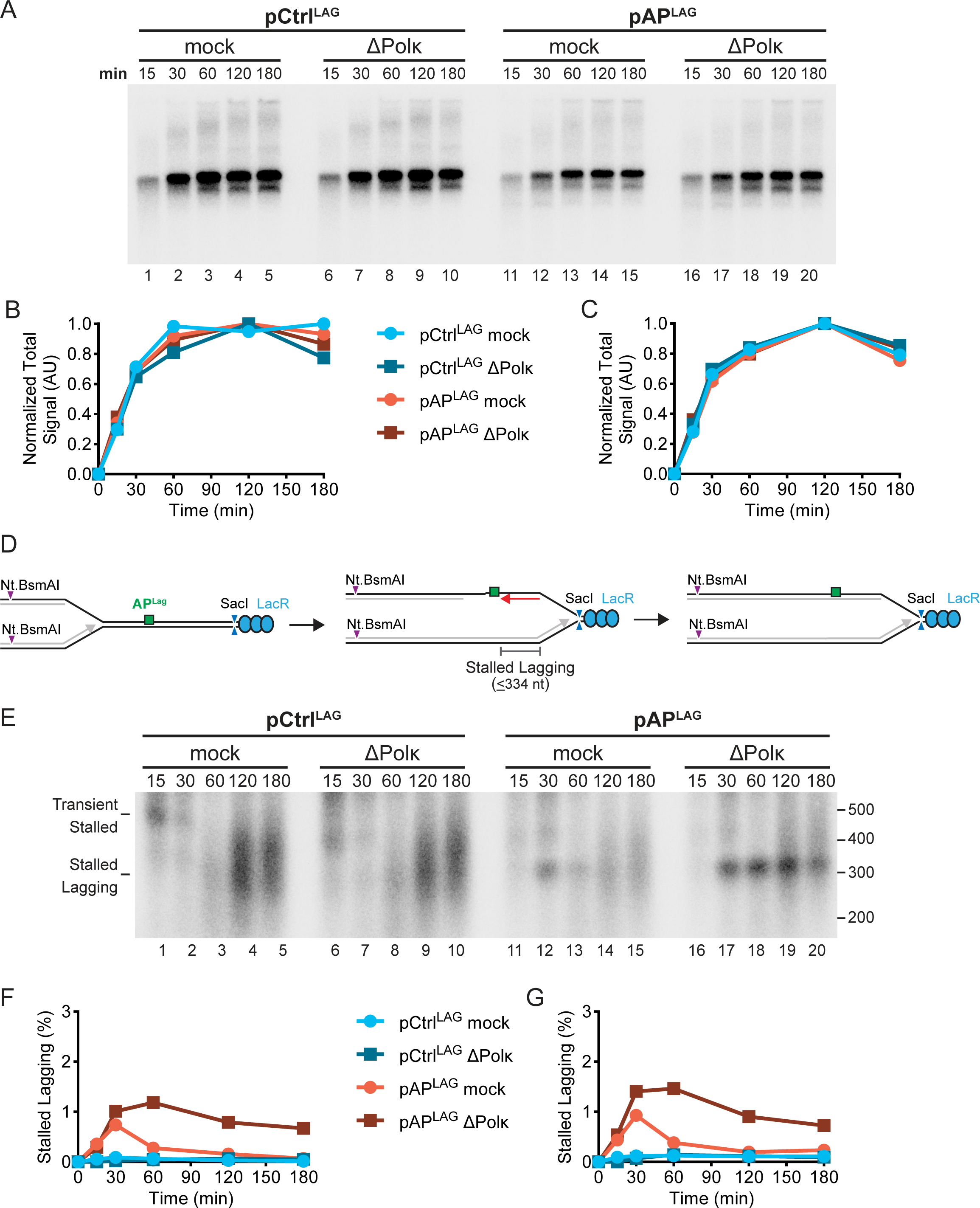
Immunodepletion of Polκ also impairs bypass of lagging strand AP sites. A) Plasmid DNA was replicated as in Fig 4A using mock- and Polκ-depleted extracts. Undigested products were separated on a native agarose gel and visualized by autoradiography. B) Quantification total synthesis from (A). Values are normalized to the peak signal recorded for each condition. C) Independent experimental replicate of (B). D) Purified products from (A) were digested with Nt.BsmAI and SacI to detect nascent lagging strands. E) Nascent strands from (D) were resolved on an alkaline denaturing agarose gel and visualized by autoradiography. F) Quantification of stalled lagging strands from (E). G) Independent experimental replicate of (F).

**Figure S13:**
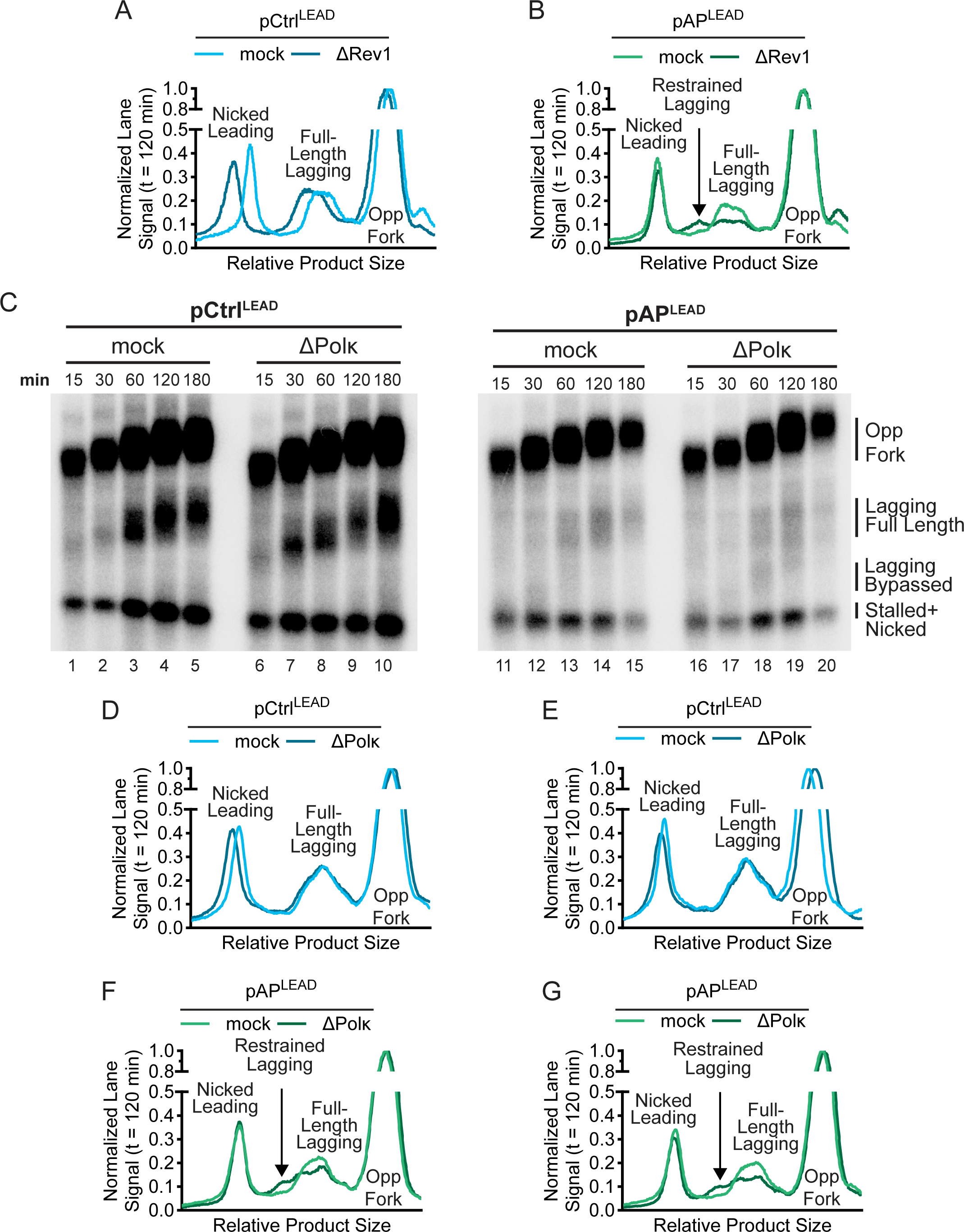
Immunodepletion of Rev1 and Polκ restrains progression of nascent lagging strands beyond the leading strand AP site. B) Independent experimental replicate of Fig. 6C. C) Independent experimental replicate of Fig. 6D. D) Plasmid DNA was replicated as in Fig 4A using mock- and Polκ-depleted extracts. Purified products were digested with Nt.BsmAI and SacI to separate nascent lagging strands. Nascent strands were resolved on an alkaline denaturing agarose gel and visualized by autoradiography. Samples for pCTRL^LEAD^ and pAP^LEAD^ were resolved in parallel using separate gels. E) Lane profiles from (C) for the control plasmid at 120 minutes. Lane signals were normalized to the peak signal of Opposing Fork (Opp Fork). F) Independent experimental replicate of (D). G) Lane profiles from (C) for the leading strand AP site plasmid at 120 minutes. Lane signals were normalized to the peak signal of Opposing Fork (Opp Fork). H) Independent experimental replicate of (F).

**Table 1.**
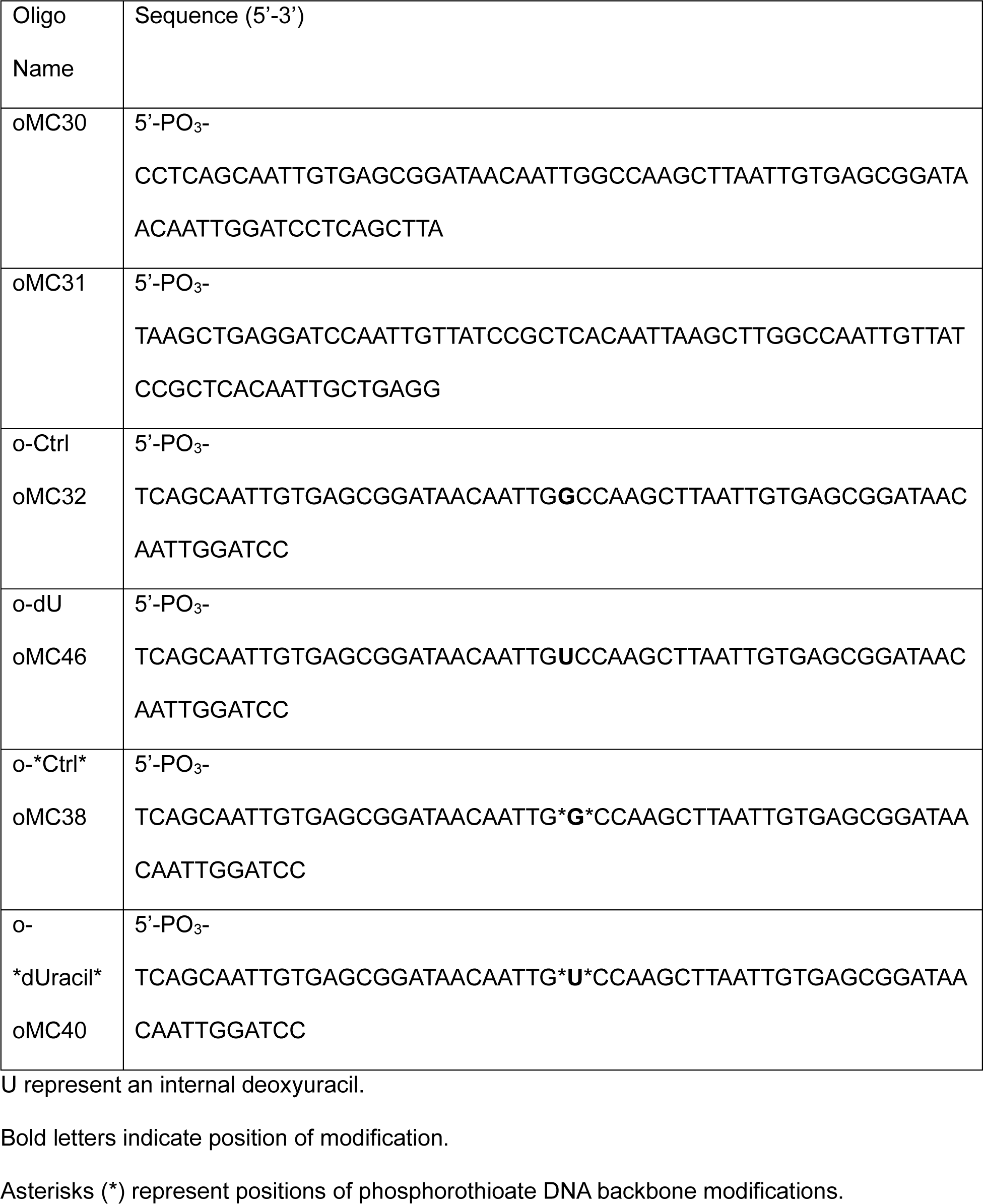
Oligonucleotides used in this study.

